# Cappable-Seq reveals the transcriptional landscape of stress responses in the bacterial endosymbiont *Wolbachia*

**DOI:** 10.1101/2024.08.15.608088

**Authors:** Youseuf Suliman, Zhiru Li, Amit Sinha, Philip D. Dyer, Catherine S. Hartley, Alistair C. Darby, Clotilde K. Carlow, Benjamin L. Makepeace

## Abstract

Bacterial endosymbionts are highly prevalent among invertebrate animals, in which they can confer fitness benefits such as pathogen defence and/or act as reproductive manipulators, inducing phenotypes including cytoplasmic incompatibility (CI). For the alpha-proteobacterium *Wolbachia*, its wide distribution among macroparasites and disease-transmitting arthropods coupled with mutualistic roles, reduction of vector competence, and CI have found recent applications in the control of several vector-borne tropical diseases. However, in common with other bacterial endosymbionts, which often lose regulatory elements during genomic erosion, the degree to which *Wolbachia* can respond to environmental or pharmacological stressors is poorly understood. Here, we apply Cappable-Seq methodology to achieve unprecedented depth and resolution of transcriptional start sites (TSS) in two *Wolbachia* strains (*w*MelPop-CLA and *w*AlbB) that have been used to transinfect mosquitoes for arbovirus control. We exposed *Wolbachia* in mosquito cell lines to temperature stress (both strains) or antibiotics (*w*AlbB only) and observed that all classes of TSS (including antisense) exhibited differential regulation, some of which were associated with mobile elements and may control ncRNA expression. Cappable-Seq also resolved the organisation of the bicistronic *cifA/cifB* operon that is responsible for inducing CI in *Wolbachia* hosts and shows great promise for revealing regulation of symbiont functions in whole invertebrates.

## INTRODUCTION

Symbiosis, the co-occurrence of dissimilar organisms in a long-term interrelationship, encompasses a highly diverse range of interactions involving every kingdom of life. Bacteria are particularly pervasive symbionts of eukaryotes and in many cases have evolved to become completely dependent on their hosts for nutrition and propagation, replicating intracellularly in an obligate dependency. In the most highly specialised symbioses, vertical transmission of the endosymbiont becomes the primary or sole mode of transfer to a new host, usually via the maternal line only (McCutcheon *et al*., 2019). As a consequence of relaxed selection and the population bottlenecks enforced by vertical transmission, genome streamlining is a key feature of inherited endosymbionts and this is accompanied by an accumulation of deleterious mutations (the so-called “Muller’s ratchet”; McCutcheon *et al*., 2019). In the most extreme cases, the bacterial genome becomes vestigial, encoding only a minimised suite of genes to support a mutualistic relationship with the host, and the distinction between endosymbiont and organelle becomes blurred (McCutcheon and Moran, 2012). However, streamlining can be slowed or even reversed by expansions of mobile elements in endosymbiont genomes; indeed, some symbiotic bacteria have higher proportions of genomic content comprised of prophages and transposons than do free-living species (Cordaux *et al*., 2008; Kaur *et al*., 2021).

A common consequence of population bottlenecks in inherited endosymbionts is a loss of DNA repair and recombination genes, which fuels the further erosion of coding capacity through pseudogenisation (McCutcheon and Moran, 2012). Consequently, the ability of an endosymbiont to regulate gene expression becomes hampered by the depletion of sigma factors and other regulatory components (Wilcox *et al*., 2003). The gamma-proteobacterial symbiont of pea aphids, *Buchnera aphidicola*, which has a genome size of 641 Kb, is a model of genome reduction in a strictly vertically-transmitted mutualist. This organism has retained only two sigma factor genes, *rpoD* (encoding housekeeping σ^70^) and *rpoH* (encoding heat-shock σ^32^), and eight transcriptional regulators (Hansen and Degnan, 2014). In experiments designed to probe the ability of *B. aphidicola* to respond at the transcriptomic level to heat stress or changes in nutritional demand due to host diet restriction, limited evidence for regulatory capacity was apparent (Reymond *et al*., 2006). However, *B. aphidicola* did regulate its transcriptome in response to host genetic background, which influences symbiont titre (Smith and Moran, 2020); although within isoclonal aphid lines, *B. aphidicola* density was affected by host rearing temperature without differential expression in the endosymbiont (Neiers *et al*., 2021). Importantly, at the protein level, differential regulation was identified in *B. aphidicola* between anatomical compartments of aphids in the absence of corresponding changes in mRNA expression (Hansen and Degnan, 2014). These protein-level differences were hypothesised to be controlled post-transcriptionally by intergenic and antisense small RNAs (sRNAs); moreover, some sRNAs were found to be differentially expressed in *B. aphidicola* when the aphid host was fed on different plants (Thairu and Hansen, 2019).

Unlike *Buchnera*, the alpha-proteobacterium *Wolbachia* has a vast host range in arthropods and some parasitic nematodes, representing the most prevalent symbiont of animals on the planet (Kaur *et al*., 2021). *Wolbachia* is of major applied significance due to its proven efficacy in controlling pest insects or reducing their ability to spread disease (Caragata *et al*., 2021), as well as representing a drug target for the treatment of filarial nematodes that cause two neglected tropical diseases : lymphatic filariasis and onchocerciasis (Johnston *et al*., 2021). Whereas *Wolbachia* in nematodes (and rare mutualistic *Wolbachia* strains in arthropods) have very small genomes in a similar range to *Buchnera* (0.5 – 1 Mb) (Lefoulon *et al*., 2020; Dudzic *et al*., 2022; Mahmood *et al*., 2023), most *Wolbachia* strains are facultative reproductive parasites of arthropods and have larger genomes (1.2 – 2.2 Mb) (Wu *et al*., 2004; Vancaester and Blaxter, 2023). These contain mobile elements such as prophages, transposons and plasmids, and recombination between *Wolbachia* strains with facultative lifestyles is possible due to uncommon but evolutionarily significant horizontal transmission events (Gomes *et al*., 2022). Of the few transcriptomic studies performed on *Wolbachia,* the majority have examined differential expression across the lifecycle of the host or between host tissues in filariae, with a recent metanalysis concluding that *Wolbachia* exhibits only a weak regulatory response involving upregulation of ribosomal proteins during larval development of the host (Chung *et al*., 2020). One study analysed the transcriptomic and proteomic response of a *Wolbachia* strain from *Drosophila melanogaster*, *w*MelPop-CLA, to doxycycline treatment in an insect cell line system (Darby *et al*., 2014). Although only 5% of genes were differentially regulated during treatment, the tolerance of *Wolbachia* to short-term therapy with tetracyclines demonstrates that this plasticity is biologically relevant, enabling a proportion of bacterial cells to survive a potent stressor. *Wolbachia* is also capable of differential expression of certain genes between host sexes (Gutzwiller *et al*., 2015); moreover, during the host lifecycle, it actively migrates from somatic to germline tissues (Fischer *et al*., 2011; Toomey *et al*., 2013). These observations suggest it responds to cues in the intracellular environment of the host to enable temporally-restricted phenotypic changes. Similarly, manipulation of the host diet in *Drosophila* affects *Wolbachia* density between somatic and germline compartments, leading to alterations in symbiont nucleoid morphology (Serbus *et al*., 2015), although transcriptional changes in *Wolbachia* in response to host diet have not been explored.

Despite major technical advances in sequencing technology over the past 15 years, transcriptomic studies of endosymbionts that cannot be cultured axenically remain challenging. This is because the abundance, and therefore sequence coverage, of host transcripts and bacterial rRNA is several orders of magnitude greater than that of endosymbiont mRNA. There are three main approaches to overcoming this problem. First, host and bacterial rRNA can be depleted using probes complementary to these hyperabundant transcripts (Kumar *et al*., 2016). This technique relies either on use of commercial kits, where the match to the rRNA species in the samples may not be optimal, or the synthesis of custom probe sets. However, even if probes have high complementarity to the target, they are limited in the scale of depletion that can be achieved. Second, terminator-5’-phosphate-dependent exonuclease (TEX) can be used to digest 5’-phosphate-associated rRNA (Sharma *et al*., 2010), although this method still lacks optimum efficacy, as the enzymatic degradation is incomplete. Finally, bait libraries, such as the Agilent SureSelect system, can be used with predesigned probes to capture rare transcripts within a mixed RNA pool (Chung *et al*., 2018). Although the Agilent SureSelect system is highly effective at enriching mappable reads, probe design can omit unique genes (*e.g*., sRNAs) that may be absent in genome annotations. It is also possible that probes could capture degraded RNA; thus, the enriched transcripts may not accurately represent the nascent transcriptional activity of the endosymbiont.

Recently, a method has been developed which involves the selective capping of prokaryotic 5’ triphosphate (5’PPP) RNA with biotinylated GTP, effectively separating primary prokaryotic mRNA from host mRNA and bacterial rRNA in the RNA pool. This method is termed Cappable-Seq (Ettwiller *et al*., 2016) and has been successfully applied to the *Wolbachia* symbiont of the filarial worm, *Brugia malayi* (Luck *et al*., 2017). This study demonstrated up to 5-fold enrichment of non-ribosomal mRNA transcripts within *Wolbachia,* allowing a more comprehensive transcriptional profiling of the endosymbiont over conventional methods. A benefit of the selective enrichment of 5’PPP is that it can define the transcriptional start site (TSS) of the captured nascent primary mRNA at single-nucleotide resolution, highlighting an early transcriptional snapshot of gene expression within the bacterium without the need for predesigned probes.

Here, using stable infections in mosquito cell lines, we apply Cappable-Seq to two *Wolbachia* endosymbionts from insects: *w*MelPop-CLA from *D. melanogaster* (McMeniman *et al*., 2008), and *w*AlbB from the tiger mosquito, *Aedes albopictus* (O’Neill *et al*., 1997). Both strains have been used in mosquito control and cause cytoplasmic incompatibility (CI), in which crosses between uninfected male hosts with infected females, or males and females infected with incompatible *Wolbachia* strains, lead to embryonic lethality (Kaur *et al*., 2021). In the context of mosquito control, CI has been used either for (a) population suppression, in which releases of incompatible males cause wild populations to crash; or (b) population replacement, in which *Wolbachia*-free wild populations are supplanted by *Wolbachia*-transinfected individuals (males and females) because the symbiont interferes with arbovirus transmission (Flores and O’Neill, 2018). An additional application of *w*AlbB has been as a surrogate for endosymbionts of filarial worms (which cannot be cultured) in antibiotic screening programmes using *Wolbachia*-infected insect cell lines (Johnston *et al*., 2014).

In this most extensive TSS study performed in an obligate intracellular bacterium performed to date, we uncover pervasive non-canonical TSS usage in both strains, including a high proportion of antisense TSS, some of which are associated with putative sRNAs. The two strains show divergent transcriptional profiles under temperature stress and differ greatly in expression of TSS linked to mobile elements (insertion sequences and prophages). Finally, in strain *w*AlbB, we reveal distinct transcriptional responses to different classes of antibiotics that have been proposed for use for filarial disease control.

## RESULTS

### Enrichment of *Wolbachia* primary transcripts

We applied Cappable-Seq to enrich for primary *Wolbachia* transcripts in strains *w*MelPop-CLA and *w*AlbB and assessed the efficacy of the process (Table 1). Processed tRNA and rRNA constituted most *Wolbachia* transcripts in unenriched samples, representing 0.21% and 8.28% of the total RNA pool, respectively; or 88.454% and 89.807% of *Wolbachia*-specific transcripts for *w*MelPop-CLA and *w*AlbB, respectively. Following enrichment, the proportion of *Wolbachia* primary transcripts increased over 94-fold (0.027% to 2.517%) and over 14-fold (0.939% to 13.535%) for *w*MelPop-CLA and *w*AlbB, respectively (Table 1). However, reads mapping to host mitochondria were also enriched 55.494 and 2.481-fold for *w*MelPop-CLA and *w*AlbB, respectively, which is expected due to the prokaryotic origin of mitochondria.

**Table 1.** Mapping rates of reads (%) to various RNA categories without (unenriched) and with (enriched) Cappable-Seq.

| RNA source | *wMelPop-CLA* |  |  | *wAlbB* |  |  |
| --- | --- | --- | --- | --- | --- | --- |
|  | Unenriched | Enriched | Fold-change | Unenriched | Enriched | Fold-change |
| Host | 99.584 | 87.144 | 0.875 | 90.736 | 85.234 | 0.939 |
| Mitochondria | 0.185 | 10.241 | 55.494 | 0.048 | 0.119 | 2.481 |
| *Wolbachia* (total) | 0.232 | 2.615 | 11.289 | 9.216 | 14.647 | 1.589 |
| *Wolbachia* tRNA | 0.057 | 0.068 | 1.206 | 3.043 | 0.194 | 0.064 |
| *Wolbachia* rRNA | 0.148 | 0.029 | 0.196 | 5.234 | 0.919 | 0.176 |
| *Wolbachia* primary transcripts | 0.027 | 2.517 | 94.134 | 0.939 | 13.535 | 14.409 |

### Identification and categorization of *Wolbachia* transcriptional start sites

We mapped 24,872,374 and 151,846,942 unique single-end 150-bp reads representing primary *Wolbachia* transcripts to their assigned TSS type (Figure 1 A) in *w*MelPop-CLA and *w*AlbB, respectively. This resolved 966 and 4,318 TSS for *w*MelPop-CLA and *w*AlbB, respectively, among all conditions tested (Figure 1, B-C). The most abundant TSS type for *w*MelPop-CLA was primary (p)TSS (42.1%), whereas for *w*AlbB, intragenic (i)TSS were predominant (45.6%). Thus, a considerable discrepancy was apparent in the total TSS detected between *w*MelPop-CLA (0.76 per kb) and *w*AlbB (2.9 per kb). However, after matching sequencing depths and TSS thresholds, the total for *w*AlbB was reduced to 0.94 per kb (Figure S1), mainly due to fewer iTSS. The least abundant TSS type for *w*MelPop-CLA and *w*AlbB were secondary (g)TSS (3.2% and 5.4%, respectively). For the annotated genes of *w*MelPop-CLA and *w*AlbB (1,286 and 1,418 respectively), approximately 29.63% (407) and 40.62% (682) harboured a canonical pTSS, respectively. Strain *w*AlbB contained a greater proportion of intragenic TSS (iTSS and asTSS), with over half of all coding sequences (CDS) containing an iTSS (53.74%), and over a third containing an asTSS (39.56%), compared to *w*MelPop-CLA (iTSS 16.10%, asTSS 10.19%) (Figure 2 A).

**Figure 1.**
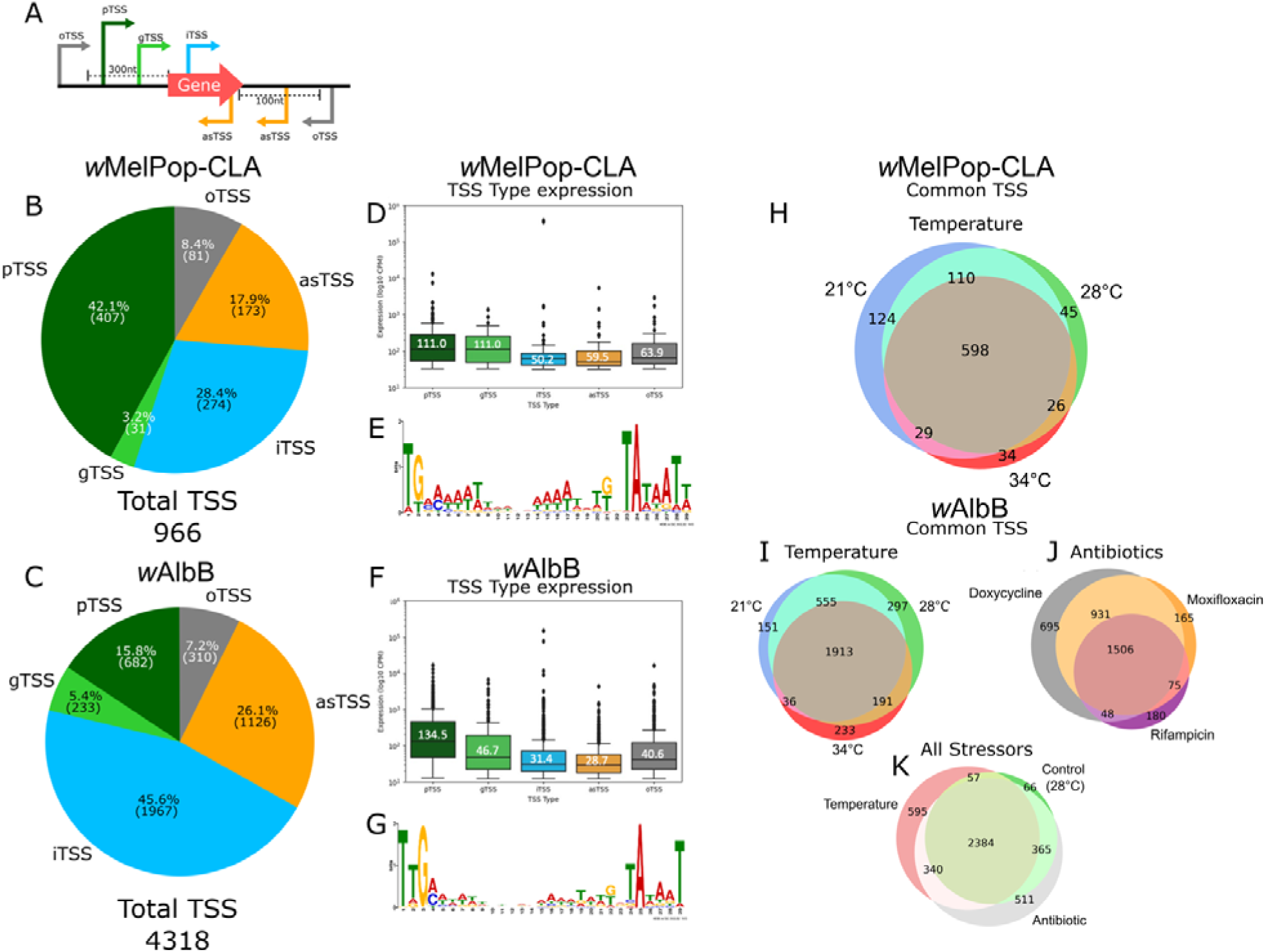
Overview of TSS characteristics for *w*MelPop-CLA and *w*AlbB. **A** Classification scheme for TSS types used in present study. Types comprise of ‘**p**TSS’, **p**rimary gene associated TSS; ‘**g**TSS’, secondary **g**ene associated TSS; ‘**i**TSS’,sense **i**nternal to an annotated gene; ‘**as**TSS’, **a**nti**s**ense to an annotated gene or within 100 nucleotides upstream antisense to the coding region; ‘**o**TSS’, and **o**rphan TSS without an associated gene. **B** Proportion of TSS types found in *w*MelPop-CLA. **C** Proportion of TSS types found in *w*AlbB. **D** TSS expression by type [median expression (log_10_ CPM)] and **E** motif from the 100 nt upstream sequence of total TSS for *w*MelPop-CLA. **F** TSS expression by type [median expression (log_10_ CPM)] and **G** motif from the 100 nt upstream sequence of total TSS for *w*AlbB. **H** Commonality of TSS between temperature stress conditions for *w*MelPop-CLA. **I** Commonality of TSS between temperature conditions for *w*AlbB. **J** Commonality of TSS between antibiotics for *w*AlbB. **K** Commonality of TSS between all stress conditions for *w*AlbB.

**Figure 2.**
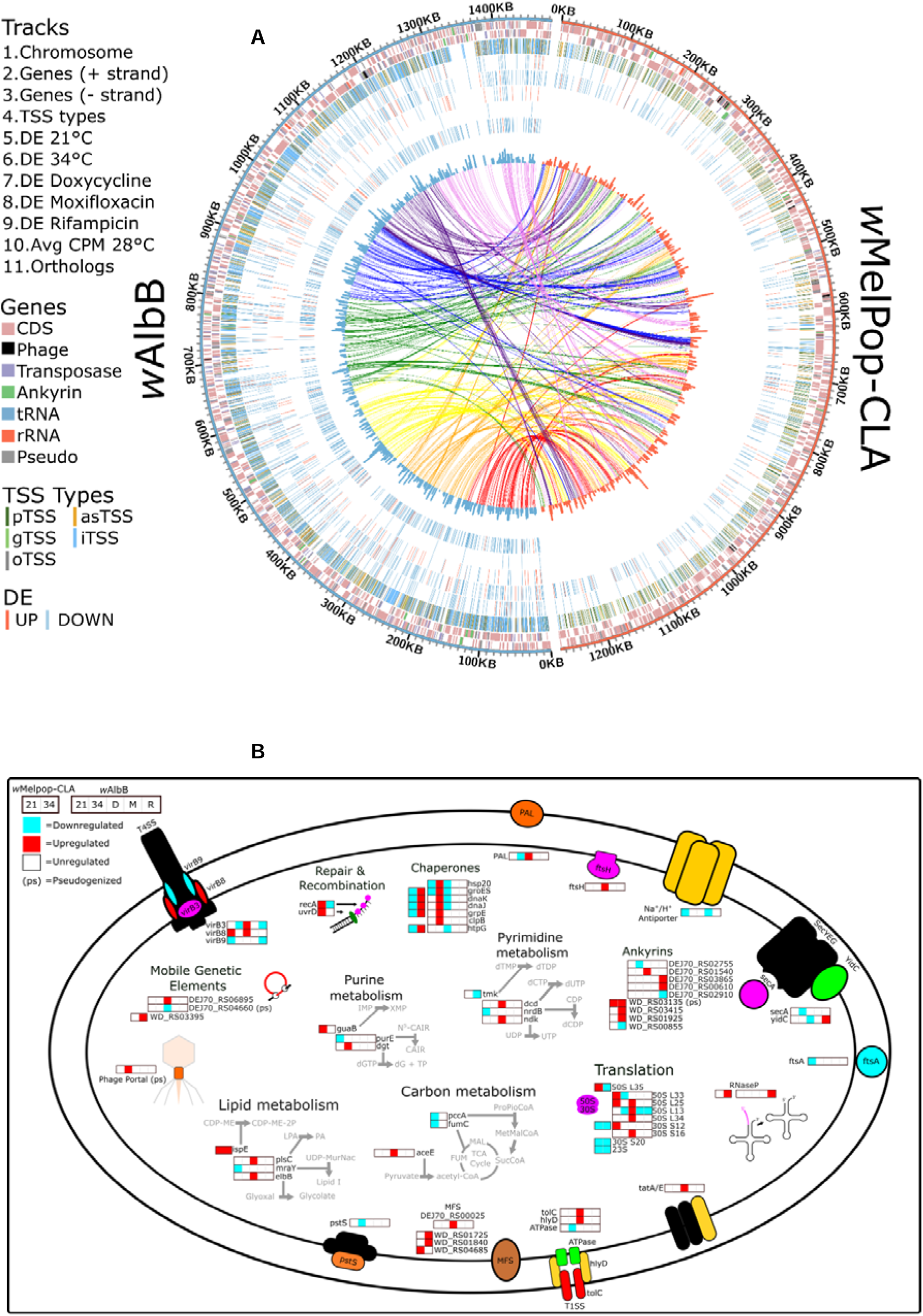
A. Genome-wide comparative mapping of gene categories, TSS distribution, DE genes, and orthologs between *w*AlbB and *w*MelPop-CLA. **B** Summary of DE pTSS and associated functional groups for both *w*MelPop-CLA and *w*AlbB under antibiotic and/or temperature stress. Metabolic pathways associated to enzymes are coloured grey. Protein structures present but not observed to be DE are filled black. Stress conditions: 21 = 21°C, 34 = 34°C, D = doxycycline, R = rifampicin, and M = moxifloxacin.

The number of TSS expressed per condition ranged from 687 - 861 and 2,373 - 2,956 under temperature stress for *w*MelPop-CLA and *w*AlbB, respectively, and 1,809 - 3,180 for *w*AlbB under antibiotic stress, with excellent concordance between biological replicates (Figure S2). Approximately one-half to two-thirds of expressed TSS were shared between temperature conditions for *w*MelPop-CLA (62%, *n* = 598) and *w*AlbB (57%, *n* = 1,913) (Figure 1, H-I); this was also apparent for *w*AlbB exposed to antibiotic stress (42%, *n* = 1,506, Figure 1 J). Conversely, for both *w*MelPop-CLA and *w*AlbB, a small proportion of TSS were expressed under specific conditions at 21°C (13%, *n* = 124 and 5%, *n* = 151), 28°C (5%, *n* = 45 and 9%, *n* = 297), and 34°C (4%, *n* = 34 and 7%, *n* = 233), respectively (Figure 1, H-I). Condition-specific TSS were also exhibited by *w*AlbB between the three antibiotic treatments [doxycycline (19%, *n* = 695), rifampicin (5%, *n* = 180) and moxifloxacin (5%, *n* = 165)]. Moreover, in *w*AlbB, a similar proportion of TSS specific for either temperature stress (14%, *n* = 595) or antibiotic stress (12%, *n* = 511) were apparent (Figure 1 K).

At baseline conditions (28°C), pTSS were the most highly expressed TSS type in both *w*MelPop-CLA and *w*AlbB, with a median expression of 111.0 and 134.5 counts per million (CPM), respectively (Figure 1, D and F). Although iTSS were highly abundant across the *Wolbachia* genomes, the median expression level for iTSS and asTSS were ranked as the lowest TSS types for both *w*MelPop-CLA and *w*AlbB. We applied motif-based sequence analysis to identify promoters, and two distinct motifs were recognized: the common AT-rich -10 TATAAT box, and a less frequent -35 TTGACA core promoter consensus sequence for σ^70^ (Figure 1, E and G).

Among the top 20 most highly expressed TSS, most TSS types were represented (excluding gTSS); 60% consisted of pTSS for both *w*MelPop-CLA and *w*AlbB (Table 2). The two most highly expressed TSS for both *w*MelPop-CLA and *w*AlbB were iTSS associated with the non-coding 6S RNA gene (*ssrS*) and the fructose-6-phospho aldolase (*fsaA*) CDS. Using a single baseline sample (28°C) for *w*MelPop-CLA and *w*AlbB, 38.3% and 8.7% of total reads were assigned to the *ssrS* iTSS (Figure S3 G and H), and 33.9% and 8.7% to the *fsaA* iTSS (Figure S3 A and B), respectively. Hypothetical proteins were also highly expressed in both strains, comprising substantial proportions of the top 20 TSS in *w*MelPop-CLA (5) and 30% in *w*AlbB (6). Notably, a pTSS for a phage tail protein (WD_RS01240) was highly abundant in the *w*MelPop-CLA dataset (Figure S3 I), as was an asTSS for a major capsid protein (WD_RS01225; Figure S3 E). Moreover, an asTSS associated to the mismatch repair protein gene *mutL* was present within the top 10 most highly expressed TSS for both *w*MelPop-CLA and *w*AlbB (Figure S3 C and D).

**Table 2.** Top 20 expressed TSS at baseline (28°C). Mean CPM represents the average expression between 28°C replicates. . Products for oTSS represent the closest annotated gene.

| wMelPop-CLA |  |  |  | wAlbB |  |  |  |
| --- | --- | --- | --- | --- | --- | --- | --- |
| TSS type | Product | Locus tag | Mean CPM | TSS type | Product | Locus tag | Mean CPM |
| iTSS | 6S RNA | WD_RS06775 | 4.20E+05 | iTSS | 6S RNA | DEJ70_RS02130 | 1.45E+05 |
| iTSS | fructose-6-phosphate aldolase | WD_RS02455 | 2.93E+05 | iTSS | fructose-6-phosphate aldolase | DEJ70_RS05135 | 3.45E+04 |
| pTSS | hypothetical protein | WD_RS02845 | 8.41E+03 | pTSS | 16S ribosomal RNA | DEJ70_RS05685 | 1.46E+04 |
| pTSS | phage tail protein | WD_RS01240 | 8.12E+03 | iTSS | signal recognition particle sRNA small type | DEJ70_RS02165 | 1.02E+04 |
| asTSS | DNA mismatch repair endonuclease MutL | WD_RS05920 | 5.39E+03 | asTSS | DNA mismatch repair endonuclease MutL | DEJ70_RS05750 | 5.96E+03 |
| iTSS | signal recognition particle sRNA small type | WD_RS06780 | 3.58E+03 | pTSS | hypothetical protein | DEJ70_RS01070 | 5.67E+03 |
| oTSS | succinate dehydrogenase assembly factor 2 | WD_RS04920 | 2.45E+03 | iTSS | 23S ribosomal RNA | DEJ70_RS05220 | 5.58E+03 |
| pTSS | 23S ribosomal RNA | WD_RS00885 | 2.25E+03 | pTSS | Rne/Rng family ribonuclease | DEJ70_RS03135 | 5.57E+03 |
| pTSS | 16S ribosomal RNA | WD_RS05540 | 2.08E+03 | pTSS | tRNA-Leu | DEJ70_RS01720 | 5.32E+03 |
| pTSS | thioredoxin family protein | WD_RS05680 | 2.04E+03 | oTSS | hypothetical protein | DEJ70_RS03120 | 5.00E+03 |
| asTSS | major capsid protein | WD_RS01225 | 1.95E+03 | pTSS | preprotein translocase subunit YajC | DEJ70_RS04650 | 4.18E+03 |
| iTSS | adenylosuccinate lyase | WD_RS03555 | 1.79E+03 | pTSS | hypothetical protein | DEJ70_RS01670 | 3.37E+03 |
| pTSS | Bax inhibitor-1/YccA family protein | WD_RS04290 | 1.76E+03 | pTSS | tRNA-Arg | DEJ70_RS04270 | 3.34E+03 |
| pTSS | hypothetical protein | WD_RS04725 | 1.64E+03 | pTSS | hypothetical protein | DEJ70_RS01345 | 3.20E+03 |
| pTSS | hypothetical protein | WD_RS00565 | 1.57E+03 | pTSS | hypothetical protein | DEJ70_RS01180 | 2.86E+03 |
| pTSS | ankyrin repeat domain-containing protein | WD_RS01925 | 1.56E+03 | pTSS | hypothetical protein | DEJ70_RS04380 | 2.78E+03 |
| oTSS | aspartate-semialdehyde dehydrogenase | WD_RS04305 | 1.23E+03 | oTSS | single-stranded-DNA-specific exonuclease RecJ | DEJ70_RS01320 | 2.77E+03 |
| pTSS | hypothetical protein | WD_RS02135 | 1.17E+03 | iTSS | tRNA-Ser | DEJ70_RS06770 | 2.65E+03 |
| pTSS | TrbC/VirB2 family protein | WD_RS02935 | 1.10E+03 | pTSS | DNA recombination protein RmuC | DEJ70_RS06410 | 2.48E+03 |
| pTSS | hypothetical protein | WD_RS00560 | 1.02E+03 | pTSS | P44/Msp2 family outer membrane protein | DEJ70_RS02180 | 2.14E+03 |

### Differential expression of primary TSS

Sequencing depth for reads mapped to *Wolbachia* were sufficient for robust DE analysis in both strains, with the exception of the 34°C condition at 48 hr for wAlbB, which was excluded from the comparisons (Figure S4). In multidimensional scaling (MDS) plots, tight clustering of biological replicates by condition and timepoint was also apparent for the vast majority of samples (Figure S5). We analysed total TSS from each strain for differential expression (DE), and of the 966 and 4,318 TSS detected, 311 (32.2%) and 688 (15.9%) TSS were found to be differentially expressed (2-fold change, FDR 5%) under at least one temperature stress condition for *w*MelPop-CLA and *w*AlbB, respectively (Table 3, Figure 2 A). The proportion of TSS types that exhibited DE reflected the available TSS repertoire. Hence, for both *w*MelPop-CLA and *w*AlbB, gTSS represented the smallest group of DE TSS, whereas the largest group was pTSS or iTSS for *w*MelPop and *w*AlbB, respectively. Under antibiotic exposure, a total of 760 DE TSS (17.6%) were detected in *w*AlbB (Table 3).

**Table 3.** Summary of *w*MelPop-CLA and *w*AlbB differentially expressed TSS for each type between all stressors. Values represent number of down- or upregulated (2-fold change, FDR 5%) TSS.

| TSS category | Wolbachia strain |  |  |  |  |  |  |
| --- | --- | --- | --- | --- | --- | --- | --- |
|  | wMelPop-CLA |  | wAlbB |  |  |  |  |
|  | No. regulated TSS per stress condition* (down, up) |  |  |  |  |  |  |
|  | 21°C | 34°C | 21°C | 34°C | Doxy. | Moxi. | Rif. |
| pTSS | 51, 16 | 135, 27 | 83, 16 | 122, 12 | 26, 40 | 2, 0 | 221, 8 |
| gTSS | 0, 0 | 6, 1 | 14, 0 | 22, 2 | 8, 2 | 2, 0 | 22, 0 |
| iTSS | 18, 6 | 65, 5 | 125, 9 | 166, 20 | 37, 26 | 19, 0 | 239, 15 |
| asTSS | 7, 3 | 35, 3 | 73, 7 | 99, 16 | 24, 15 | 7, 0 | 89, 27 |
| oTSS | 4, 5 | 14, 6 | 18, 13 | 22, 8 | 9, 3 | 1, 0 | 30, 8 |
| Total | 80, 30 | 255, 42 | 313, 45 | 431, 58 | 104, 86 | 31, 0 | 601, 58 |

\*Doxy., doxycycline; Moxi., moxifloxacin; Rif., rifampicin.

Of the 175 *w*MelPop-CLA DE pTSS under temperature stress, 16 and 27 were upregulated at 21°C and 34°C, respectively, whereas of the 189 *w*AlbB DE pTSS, 16 and 12 were upregulated at 21°C and 34°C, respectively (Table 3). For both *w*MelPop-CLA and *w*AlbB, the majority of DE pTSS were downregulated in response to temperature stress (Table 3, Figure 2 A). Of the 760 *w*AlbB DE pTSS identified in response to antibiotic stress, 40, 0, and 8 were upregulated; while 26, 2, and 221 were downregulated under doxycycline, moxifloxacin, and rifampicin stress, respectively (Table 3). Thus, in common with temperature stress, rifampicin exposure mostly induced downregulation of pTSS, whereas doxycycline exposure led to a greater proportion of upregulated pTSS (Figure 2 A). Surprisingly, the TSS expression patterns under moxifloxacin exposure were quite similar to control conditions (Figure 2 A), as reflected in the MDS plots (Figure S5 C).

To determine whether certain functional categories were overrepresented in the DE datasets, we applied KEGG pathway mapping. In both *w*MelPop-CLA and *w*AlbB, DE pTSS were associated with chaperones and heat-shock proteins (HSPs), translation, repair and recombination, nucleotide metabolism, lipid metabolism, mobile genetic elements, ankyrins, secretion and transport, and carbon metabolism (Figure 2 B, Figure 3). The largest group of upregulated pTSS during temperature stress were chaperones and HSPs for both *w*MelPop-CLA and *w*AlbB (Figure 2 B, Figure 3). Remarkably, pTSS associated with a phage portal protein pseudogene (DEJ70_RS00855) or transposase (WD_RS03395) were upregulated at 34°C for *w*AlbB and *w*MelPop-CLA, respectively (Figure 2 B). Primary TSS proximal to other mobile elements in *w*AlbB were affected by doxycycline treatment, as evidenced by upregulation of a transposase (DEJ70_RS06895), alongside downregulation of an apparent pseudogenised transposase (DEJ70_RS04660; Figure 2 B). Among antibiotic stressors, doxycycline induced the largest category of upregulated pTSS involved in secretion and transport (Figure 3 A).

**Figure 3.**
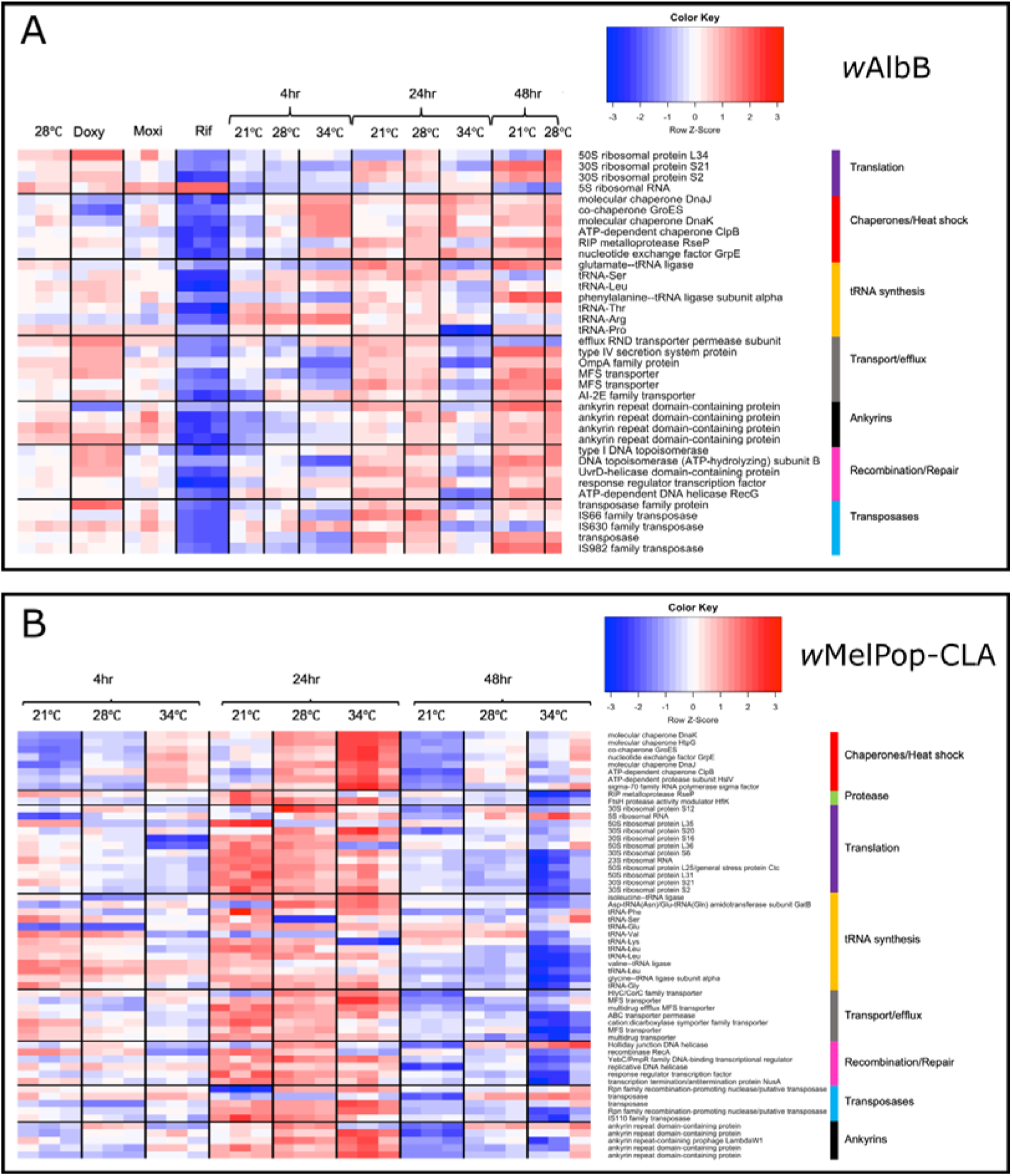
Summary heatmap of DE pTSS for **A** *w*AlbB and **B** *w*MelPop-CLA. Associated functional groups are displayed on right. Expression levels represent Z-scores (sample CPM minus mean CPM of associated row, divided by SD of CPMs in that row).

We identified orthologous genes shared by *w*MelPop-CLA and *w*AlbB pTSS (Figure 2 A) and analysed DE between pTSS of each strain. Forty pTSS were classed as DE under at least one temperature stress condition (Figure 4); these were located upstream of genes encoding heat shock proteins (*dnaJ, groES, dnaK, dnaJ, htpG*), proteases (*clpB*), T4SS secretion components (*virB9*), translation (ribosomal proteins), cell division (*ftsZ*), outer membrane proteins (*ompA*), and hypotheticals. Both strains also exhibited regulation of heat shock-related protease genes such as *hslV* and *clpB*; HSPs displayed upregulation at 34°C and downregulation at 21°C. In contrast with *w*MelPop-CLA, *w*AlbB did not display DE of the pTSS for *rpoH* (encoding σ^32^).

### Putative small RNAs associated with antisense transcriptional start sites

To determine if asTSS may be associated with sRNA expression, the top 20 asTSS in *w*MelPop-CLA were sorted according to their average expression (Figure 5C), which ranged from 49 to 5,389 CPM. Minimum free energy (MFE) values, which are inversely related to the stability of RNA secondary structures, were calculated for sRNAs predicted downstream of asTSS. These ranged from -7.5 to -30.7 at 28°C, compared with a mean MFE of ∼-14 for randomised 100-bp sequences from both *Wolbachia* strains. Seventeen of the top 20 asTSS from wMelPop-CLA displayed evidence of DE: *WsnRNA-59,* associated with a phage major capsid gene (Figure 5 C, rank 4), was upregulated over 5-fold at 21°C and downregulated at 34°C, while other asTSS that may target a major capsid protein gene (WD_RS02715; Figure 5 C, rank 17) or *rpoB* (Figure 5 C, rank 6) were downregulated more than two-fold under 34°C exposure. Of the top 20 asTSS present in *w*AlbB, the average expression ranged from 113 to 5,962 CPM, whilst MFE values encompassed -5.9 to -26.2 (Figure 5 D). Unlike *w*MelPop-CLA, asTSS related to prophage genes were not prominently expressed in *w*AlbB, although one was associated with a major capsid pseudogene (Figure S3 F). Evidence for DE was observed for 11 of the top 20 *w*AlbB asTSS under at least one stressor, especially for rifampicin exposure, which resulted in upregulation of six asTSS. An asTSS in opposite orientation to the mismatch repair protein gene *mutL* was the most highly expressed asTSS for both *w*MelPop-CLA (WD_RS05920) and *w*AlbB (Figure S3 C and D), with an average expression of 5,389 and 5,962 CPM, respectively, at 28°C. Both of the asTSS for *mutL* displayed DE, exhibiting downregulation at either 34°C (*w*MelPop-CLA) or 21°C (*w*AlbB). Unusually for Pseudomonadota, many *Wolbachia* genomes harbour two *mutL* paralogues (Wu *et al*., 2004), one of which is located in the eukaryotic association module of phage WO in strains containing prophages (Bordenstein and Bordenstein, 2016), such as *w*Mel and its derivatives. However, the asTSS was proximate to the *mutL* paralogue positioned outside prophage regions. In both *w*MelPop-CLA and *w*AlbB, RNA secondary structures with low MFE were predicted within the antisense sequence of the *mutL* gene, and the asTSS associated with *w*AlbB *mutL* was expressed at a 43-fold higher level than the pTSS in the sense orientation. Remarkably, the pTSS for the other *mutL* paralogue in *w*AlbB was poorly expressed, whereas in *w*MelPop-CLA, neither *mutL* paralogue had a detectable pTSS.

**Figure 4.**
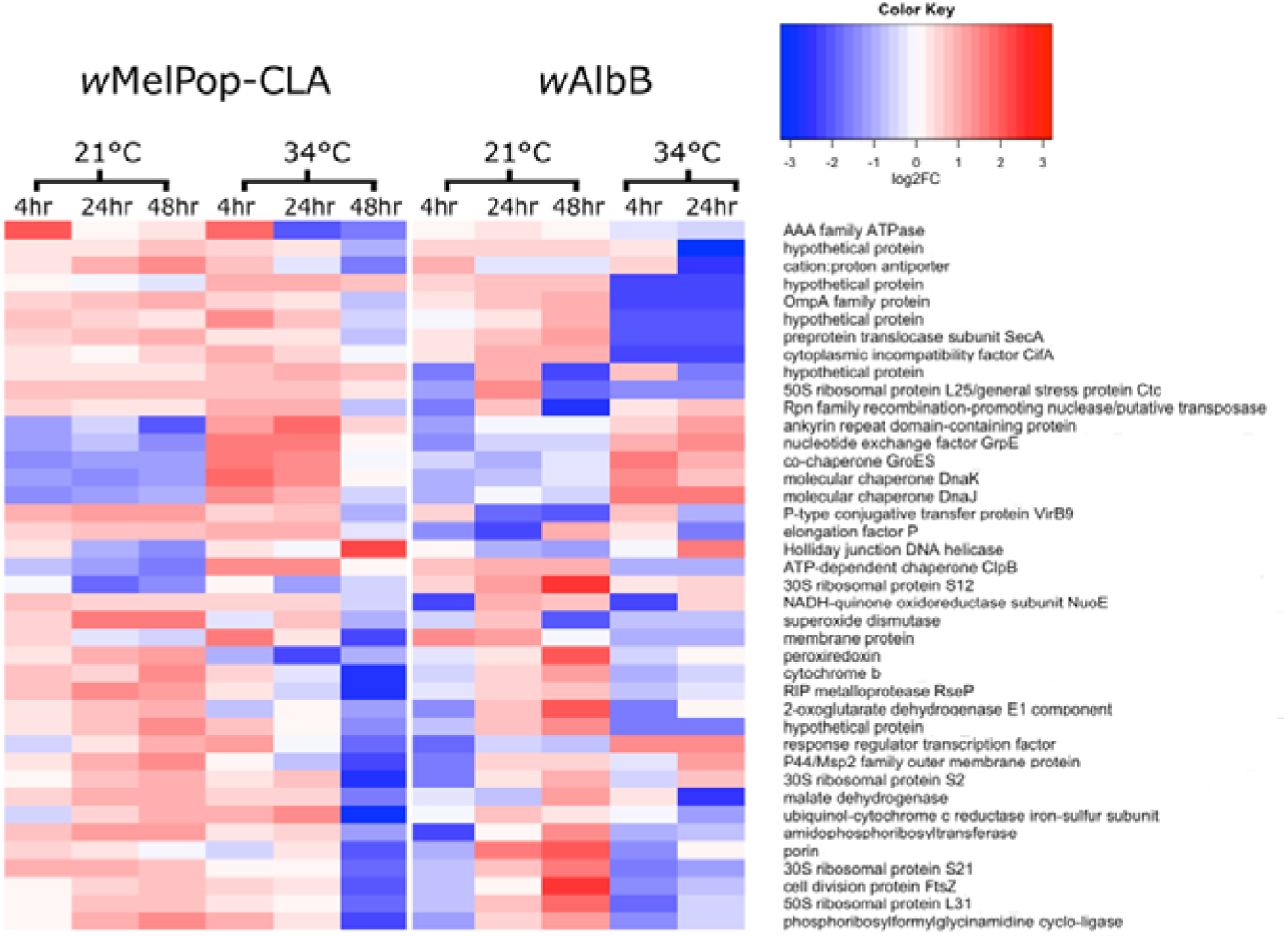
Heatmap of differentially expressed orthologous pTSS exposed to temperature stress. Orthologous pTSS were located upstream of genes encoding heat shock proteins, ankyrins, secretion systems, and hypothetical proteins. The colour key is shared between strains, but the normalized value is specific to the individual strain. Expression levels displayed in Z-scores (sample CPM minus mean CPM of associated row, divided by SD of CPMs of that row within each strain).

**Figure 5.**
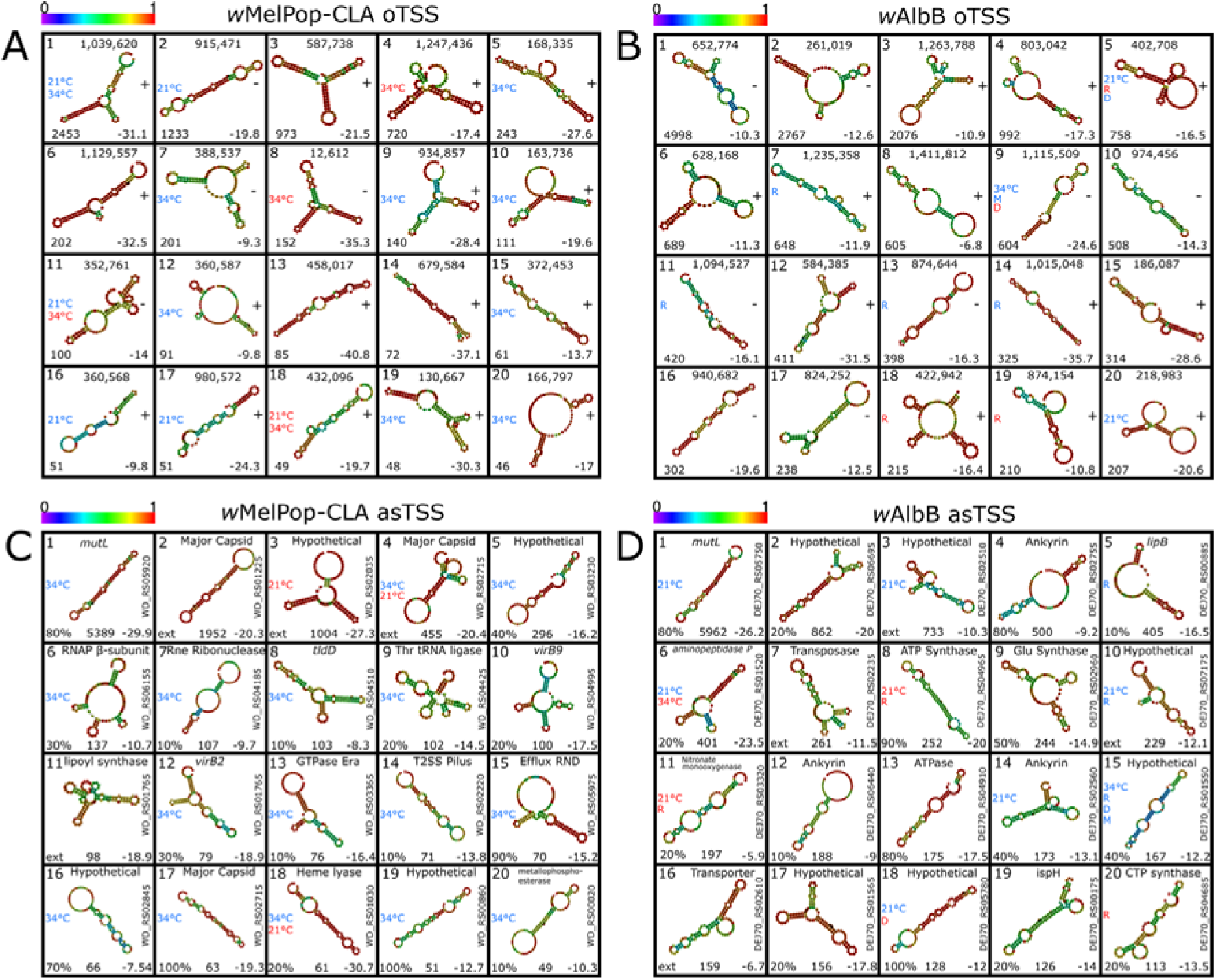
Summary of top 20 ncRNA and associated characteristics. Top 20 expressed oTSS for **A** *w*MelPop-CLA and **B** *w*AlbB. Top 20 expressed asTSS for **C** *w*MelPop-CLA and **D** *w*AlbB. Centre, predicted RNA secondary structure; top left, rank by expression; top centre, genome coordinate for oTSS or target gene for asTSS; right, strand for oTSS or locus for asTSS; bottom right, MFE value; bottom centre, average CPM for asTSS; bottom left, average CPM for oTSS or location along the gene as a percentage for asTSS; left, stressors which induced DE. Stressors are labelled R, rifampicin; M, moxifloxacin; D, doxycycline; 21°C and 34°C. Red or blue coloured text indicates up- or downregulation, respectively. Colour key spectrum above each panel represents base-pair probabilities.

### Highly expressed intergenic and orphan transcriptional start sites

An iTSS positioned within the *fsa* gene was consistently in the top two most highly expressed TSS for both *w*MelPop-CLA and *w*AlbB under all tested conditions but showed no evidence of DE (Figure S3 A and B). The location of the *fsa* iTSS was identical for both *w*MelPop-CLA and *w*AlbB, suggesting high conservation and a common function. The *fsa* gene encodes fructose-6-phosphate aldolase, which catalyses the reversible formation of fructose 6-phosphate (F6P) from glyceraldehyde 3-phosphate and sedoheptulose 7-phosphate (Schürmann and Sprenger, 2001). However, for F6P to be utilized for glycolysis requires phosphorylation to fructose 1,6-bisphosphate by phosphofructokinase, which is absent in *Wolbachia*. Moreover, the highly expressed iTSS of *fsa* was located downstream from the active site of the enzyme, such that if the transcript was translated into protein, it would be incapable of contributing to F6P synthesis. The RNA sequence 100 nt downstream of the *fsa* iTSS exhibited a secondary structure with low MFE in both *w*MelPop-CLA and *w*AlbB (Figure S1 A and B), suggesting an alternative role as an ncRNA.

Of the 81 oTSS detected for *w*MelPop-CLA, the most highly expressed included three associated with *Wolbachia* ncRNAs predicted in an earlier study on *w*Mel (Woolfit *et al*., 2015). The oTSS at position 1,039,620 (Figure 5 A, rank 1) was identified as *ncrwMel02* and was the most strongly expressed oTSS in this strain; moreover, the transcript was downregulated during exposure to temperatures of 21°C or 34°C. The small ncRNA designated *ncrwMel01* was detected at genomic position 587,738 (Figure 5 A, rank 3), while the third oTSS associated with a previously-identified ncRNA from *w*Mel was located at position 67,860 (-strand) but exhibited very low expression. Whereas neither of these transcripts displayed evidence of DE, an oTSS located at genomic position 915,471 (Figure 5 A, rank 2) - incidentally not identified in the study of Woolfit *et al*. (2015) - showed downregulation at 21°C. In contrast, among the 310 oTSS discovered in *w*AlbB, very few highly-expressed intergenic ncRNA candidates were regulated under thermal stress (3/20 compared with 16/20 in *w*MelPop-CLA; Figure 5 A and B).

### Predicted operon structures

We applied operon prediction to the genomes of both *Wolbachia* strains using TSS data. Of the 1,294 and 1,406 genes in *w*MelPop-CLA and *w*AlbB, respectively, 233 (18%) and 212 (15%) were predicted to be arranged into operons of two or more genes (Tables S1 and S2). The majority of operon-associated genes were predicted to be situated in gene pairs, representing 70% of operons in both *w*MelPop-CLA and *w*AlbB. The largest predicted operons were associated with the 30/50S ribosomal proteins, which contained 28 and 23 genes for *w*MelPop-CLA and *w*AlbB, respectively. Operons with a robustly located pTSS accounted for 17% (*n* = 40) of the repertoire in *w*MelPop-CLA compared with 27% (*n* = 58) in *w*AlbB. The least frequent TSS type associated with predicted operons was alternate gTSS, which accounted for just two and ten operons in *w*MelPop-CLA and *w*AlbB, respectively.

The T4SS operons are of particular interest in *Wolbachia* biology due to the potential importance of secreted effectors in manipulating host cells, as well as the DE exhibited by some of its components for *w*AlbB, but not *w*MelPop-CLA, under temperature or antibiotic stress (Figure 2 B; Darby *et al*., 2014). In both strains, the first operon (T4SS-1) is proximal to a surface protein gene (Rancès *et al*., 2008), *wspB*, pseudogenisation of which is associated with thermotolerance in natural genomic variants of *w*Mel and *w*AlbB strains (Hague *et al*., 2021; Gu *et al*., 2022; Martinez *et al*., 2022). We inspected the local region of the T4SS-1 operon, which contains eight genes from *tRNA^Leu^* – *wspB* in both *w*MelPop-CLA and *w*AlbB (Figure 6), and located a pTSS approximately 37 nucleotides upstream of the initiation codon of *tRNA^Leu^*. However, the TSS landscape across the operon displayed striking differences between the strains. In T4SS-1 of *w*MelPop-CLA, the operon contained a total of eight TSS, comprising two pTSS, two iTSS, and four asTSS. The second pTSS for *w*MelPop-CLA was located in the intergenic region between *virD4* and *wspB*, approximately 20 nucleotides upstream of the *wspB* initiation codon. In contrast, the *w*AlbB-CLA T4SS-1 operon contained a total of 23 TSS, comprising three pTSS, eleven iTSS, eight asTSS, and one pseudogene-associated pTSS (pTSSps). The three pTSS associated with the *w*AlbB T4SS-1 operon were upstream to *tRNA^Leu^*, *virB8*, or *virD4*; whereas the pTSSps was 20 nucleotides upstream of the initiation codon for the *wspB* pseudogene.

**Figure 6.**
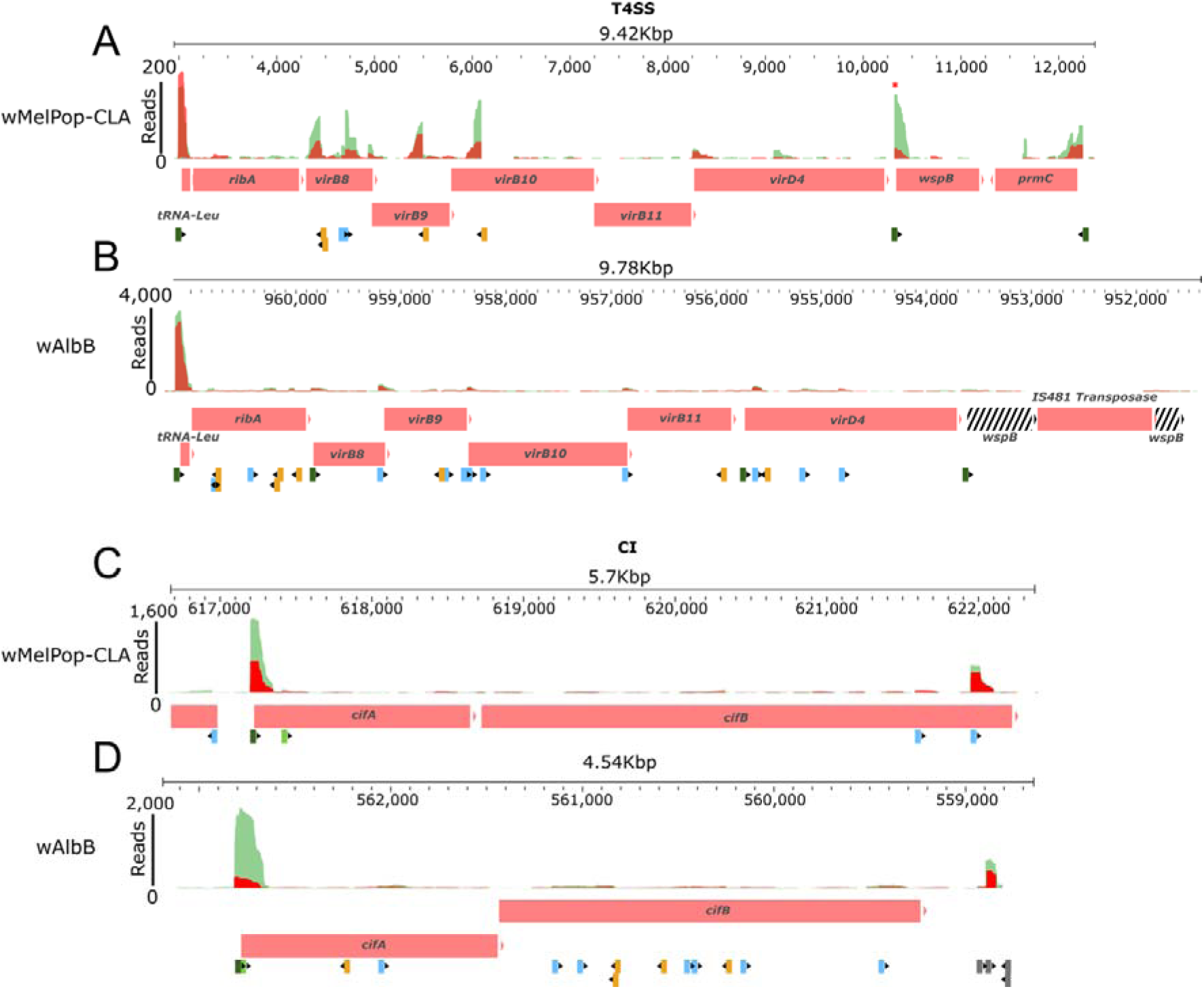
Overlay of TSS in relation to associated operon. The T4SS-1 operon for **A** *w*MelPop-CLA and **B** *w*AlbB. The CI operon for **C** *w*MelPop-CLA and **D** *w*AlbB. Genes are coloured as red boxes; pseudogenes are black and white striped. Green and red peaks represent mapped reads under 28°C and 34°C temperatures, respectively. The TSS are indicated below the associated operon and coloured per TSS type as dark green, pTSS; light green, gTSS; light blue, iTSS; orange, asTSS; grey, oTSS. Red asterisk highlights downregulation of *w*MelPop-CLA *wspB* during heat stress (34°C).

We also examined the local region of the *cifA/cifB* CI operon, as limited data are currently available for its regulation in any strain. This revealed a single pTSS that was 25 and 32 nucleotides upstream of the *cifA* initiation codon in *w*MelPop-CLA and *w*AlbB, respectively, indicating a shared bicistronic arrangement (Figure 6). The *w*MelPop-CLA CI operon contained four TSS, comprising one pTSS, one gTSS, and two iTSS; whereas the *w*AlbB CI operon contained 14 TSS (five asTSS in addition to one pTSS, one gTSS, and seven iTSS). The *cifA* pTSS in *w*AlbB exhibited downregulation at 34°C, while one of the asTSS associated with *cifB* displayed upregulation under rifampicin exposure, albeit from a very low baseline.

## DISCUSSION

Prior to our study on *Wolbachia*, few studies on the primary transcriptome of obligate intracellular or other uncultivable bacteria had been performed. We previously applied a TEX-based protocol to *w*MelPop-CLA under doxycycline stress but did not analyse TSS distribution in that experiment (Darby *et al*., 2014). Previous studies on *Chlamydia* spp. and *Candidatus* Phytoplasma asteris using the TEX method have identified several hundred TSS, equating to approximately 0.3 – 0.4 per kb (Albrecht *et al*., 2009, 2011; Nijo *et al*., 2017). This was surpassed by a TEX-based analysis of *Rickettsia conorii* TSS expression in tick and mammalian cell lines (Narra *et al*., 2020), which uncovered several thousand TSS (∼3 per kb), almost identical to what we attained here for *w*AlbB. However, our study on *Wolbachia* was unique in applying replicated experiments across several stressors in two different strains. Indeed, a replicated Cappable-Seq study involving DE analysis of TSS across multiple stress conditions was achieved only recently in *E. coli,* which identified 19,975 TSS (3.6 per kb) but was limited to a single strain (Zehentner *et al*., 2023).

The iTSS category was consistently the most poorly expressed type within both strains, suggesting that deeper sequencing depth preferentially uncovers more iTSS over other TSS classes. This may place limits on the extent to which deeper sequencing can uncover biologically relevant TSS. However, in *E. coli* - as here with *Wolbachia* - iTSS were consistently detected across biological replicates and many exhibited DE between different conditions (Zehentner *et al*., 2023). Thus, the total count of TSS for *w*MelPop-CLA would have been expected to be larger if the number of stress-inducing conditions was equivalent to those applied to *w*AlbB. While canonically, only a single origin of initiation upstream to the coding sequence of a gene is expected (*i.e*., a pTSS), the pervasiveness of iTSS we observed here is not limited to *E. coli*; other bacteria exhibiting high proportions of iTSS include *Mycobacterium marinum* (38%) (Huang *et al*., 2021), *Salmonella enterica* (11%) (Ramachandran, Shearer and Thompson, 2014), and *Bacteroides thetaiotaomicron* (45%) (Ryan *et al*., 2020).

For both *w*MelPop-CLA and *w*AlbB, the 6S RNA (encoded by *ssrS*) was consistently amongst the most highly expressed TSS. This ncRNA is known to form a hairpin secondary structure in which a pre-melted bubble mimics a DNA promoter capable of binding to the RNAP holoenzyme (Eσ). In *E. coli,* the bound 6S-Eσ complex results in downregulation of σ^70^ regulated genes by competitive binding to RNAP (Wassarman and Storz, 2000). The expression of 6S RNA has previously been detected in both filarial and arthropod *Wolbachia* strains. Darby *et al*. (2012) observed upregulation of 6S RNA from strain *w*Oo in the gonads of the filarial nematode *Onchocerca ochengi* relative to somatic tissue, whereas Darby *et al*. (2014) reported downregulation of 6S RNA in *w*MelPop-CLA strain exposed to doxycycline. The 6S rRNA of *w*AlbB was also shown to be upregulated in response to co-infection of mosquito cells by dengue virus *in vitro* (Leitner *et al*., 2021), as well as in *R. conorii* during an infection time-course in mammalian cells but not tick cells (Narra *et al*., 2020). During the lifecycle of the filarial worm *Brugia malayi*, expression levels of 6S RNA in *Wolbachia* strain *w*Bm correlated with symbiont replication rates (Chung *et al*., 2019). We did not observe regulation of *ssrS* in the current study, probably because 6S RNA acts as a template for product RNAs that are capped *in vivo* (Bonar *et al*., 2022), and these processed transcripts would not be captured during the initial stage of the Cappable-Seq protocol. Few functional studies of 6S RNA have been performed in intracellular bacteria, but deletion of *ssrS* in *Legionella pneumophila* resulted in a 10-fold reduction in intracellular growth along with downregulation of genes involved in amino acid metabolism and stress adaptation such as *groES* (Faucher *et al*., 2010). The 6S RNA is also essential for an effective oxidative stress response in *E. coli* (Burenina *et al*., 2022). The very high abundance of 6S RNA observed in the current study suggests it not only controls intracellular replication but also prioritizes expression of stress-related genes, such as *groES,* with less reliance on sigma factors. Indeed, we only detected the presence of a sequence motif that resembles the -35 (TTGACA) and -10 (TATAAT) boxes of *E. coli* σ^70^, despite expression of σ^32^ in both *Wolbachia* strains and signs of its regulation in *w*MelPop-CLA, although this failure could be due to divergence of promoter motifs in the symbiont compared to free-living bacteria. *Wolbachia* genomes may also encode unidentified extracytoplasmic functional sigma factors, as has been suggested for *Rickettsia* spp. (Narra *et al*., 2020).

Numerous studies have demonstrated that *Wolbachia* strains display marked differences in susceptibility to heat stress. Following early trials with the virulent strain *w*MelPop and its cell culture-adapted derivative, *w*MelPop-CLA, most field releases of *Wolbachia* for population replacement of *Aedes aegypti* worldwide have utilised *w*Mel due to its low fitness costs (Walker *et al*., 2011), although *w*AlbB is also well-tolerated by *Ae. aegypti* and has been the strain of choice for the Malaysian dengue control programme (Nazni *et al*., 2019). Both *w*MelPop-CLA and *w*Mel are sensitive to environmentally-relevant heat stress, such as cycling of larval development temperatures from 26-37°C (Ross *et al*., 2017). Under laboratory conditions, this led to reduced symbiont density and loss of the CI phenotype for *w*MelPop-CLA, whereas *w*AlbB was significantly more tolerant and retained the ability to induce complete CI. However, data from the field after releases in Cairns, Queensland, showed that *w*Mel frequencies in transinfected mosquitoes can recover following a heatwave that reached 43.6°C (Ross *et al*., 2020).

After being exposed to 34°C heat stress for four hours, both *w*MelPop-CLA and *w*AlbB upregulated HSP genes from the GroESL and KJE chaperone systems as expected. However, differential expression of *rpoH* under thermal stress was only apparent in *w*MelPop-CLA. Thus, *w*AlbB appears to upregulate HSP genes without, or with minimal contribution from, σ^32^. The heat-shock response peaked at 24 h and by 48 h, *w*MelPop-CLA exhibited mass downregulation of all pTSS, including HSP genes, which may be indicative of cell death. Upregulation of *Wolbachia* HSPs during changes in the host cell environment have been observed previously in response to DENV infection (Leitner and Bishop, 2021) and host development (Grote *et al*., 2017; Chung *et al*., 2019). Following cold stress at 21°C, downregulation of GroESL or KJE components was apparent in *w*AlbB and *w*MelPop-CLA, respectively, reflecting an additional similarity in the two strains’ responses to temperature stress. Notably, it has been demonstrated *w*Mel has co-adapted with *D. melanogaster* along latitudinal temperature clines in Australia and other continents, with a recently-derived tropical variant exhibiting lower maternal transmission fidelity at cool temperatures (down to 20°C) than does a temperate variant (Hague *et al*., 2022). Unlike *w*MelPop-CLA in which the *wspB* gene is intact, tropical variants of *w*Mel are associated with a premature stop codon at this locus, and strain wMelM maintained increased resilience to heat stress when transferred into *Ae. aegypti* (Gu *et al*., 2022). We found a *wspB* pTSS that is independently regulated of the T4SS-1 operon of wMelPop-CLA; this alternative transcriptional unit (ATU) was downregulated at 34°C. Similar to *w*Mel, two variants of *w*AlbB exist in the wild and exhibit alterations in thermal sensitivity that are associated with a polymorphism in *wspB* among other genomic differences (Martinez *et al*., 2022), with the *wspB-*disrupted variant being used in culture here. Our data suggest that the pseudogenisation of the *wspB* in *w*AlbB is accompanied by loss of regulation under temperature stress, although a remnant pTSS is associated with this disrupted ORF.

Despite their reduced genomes, *Wolbachia* strains from arthropods harbour an unusually high number of insertion sequences (IS), representing 6% or 13% of genomic content for *w*Mel and *w*AlbB, respectively (Wu *et al*., 2004; Sinha *et al*., 2019). Intact or degraded prophages originating from phage WO are also common in the arthropod symbionts. In *w*MelPop-CLA, we observed upregulation of a transposase gene at 34°C, which progressively increased expression throughout exposure to elevated temperature, whereas doxycycline treatment induced upregulation of IS genes in *w*AlbB. Hence, similarly to phage WO in *Wolbachia*-infected *Nasonia vitripennis* wasps (Bordenstein and Bordenstein, 2011), some IS elements may be responsive to temperature increases, while upregulation of transposases as a result of antibiotic treatment has also been reported in free-living bacteria (Schreiber *et al*., 2013). However, while the *w*MelPop-CLA genome contains two main prophage regions, no evidence of entry into a lytic cycle during heat stress was apparent here, as only a single phage structural component (phage tail protein, WD_RS02845) showed high expression from a pTSS. Instead, several asTSS associated with phage major capsid proteins were among the top-20 highly expressed asTSS in wMelPop-CLA.

Transcripts antisense to major capsid protein genes in the WO-A and WO-B prophage regions were previously observed in RNA-Seq studies on the susceptibility of *w*MelPop-CLA to doxycycline (Darby *et al*., 2014) and in *w*Mel during progression through the *D. melanogaster* lifecycle (Gutzwiller *et al*., 2015). The antisense transcript overlapping the major capsid protein E gene in the WO-B prophage region was designated *WsnRNA-59* in a study of *Wolbachia*-host interactions (Darby *et al*., 2014; Mayoral *et al*., 2014). The presence of these antisense transcripts to phage capsid protein genes suggests that *w*MelPop-CLA may be suppressing activation of the lytic cycle of phage WO via antisense-sense complementary degradation. The use of asRNA to inhibit the lytic cycle of phage has been observed in P22, hosted by *Salmonella* Typhimurium, which is repressed by the *c*_2_ protein that blocks transcription of proteins required for lytic cycle development (Ballivet and Eisen, 1978). Lytic growth of the P22 phage is activated by its anti-repressor protein (Ant) that counteracts *c*_2_ repression. However, a phage-encoded asRNA, Sar, can prevent the lytic cycle by blocking the *ant* ribosome binding site, inhibiting translation of the anti-repressor (Schaefer and McClure, 1997). *Salmonella* Typhimurium also expresses an asRNA, Isrg, levels of which were inversely correlated with expression of a phage tail component (Padalon-Brauch *et al*., 2008). Accordingly, the presence of genes encoding the dsRNA-specific RNase III in both *w*MelPop-CLA and *w*AlbB is consistent with a potential gene-silencing mechanism. In addition to phage-related asRNA, several transposase genes were associated with asTSS in both *Wolbachia* strains. Recently, antisense transcription has been hypothesized to suppress expression of Rickettsiales-amplified genetic elements (RAGE) at the protein level in *Orientia tsutsugamushi*, a pathogen in the same order as *Wolbachia* (Rickettsiales), since RAGE products with high ratios of antisense to sense transcription were not detectable by proteomics in contrast with those exhibiting low ratios (Mika-Gospodorz *et al*., 2020).

Phage portal proteins are key initiators of capsid assembly (Cuervo and Carrascosa, 2012); therefore, the DE associated with portal pseudogenes observed here may reflect regulatory remnants of previously active phages in the evolutionary history of *Wolbachia* that have yet to be completely lost. Pervasive transcription from pseudogenes (not limited to mobile elements) has also been reported from bacteria with more extensively degraded genomes than those of arthropod *Wolbachia*, such as *Mycobacterium leprae* and the tsetse fly symbiont, *Sodalis glossinidius*. In *M. leprae*, some pseudogenes have retained functional promoters (Williams *et al*., 2009), while in *S. glossinidius,* DE involving pseudogenes has been observed (Goodhead *et al*., 2020). Thus, expression from a pTSS for the *wspB* pseudogene in *w*AlbB indicates that it may have a regulatory role associated with the T4SS-1 operon.

The identity of some of the most highly-expressed alternative TSS was surprising, such the asTSS associated with a *mutL* paralogue, which was the dominant antisense transcript in both strains. When interpreted in the context of poor *mutL* pTSS expression, this suggests suppression of mismatch repair in *Wolbachia*. Mutations in *mutL* have been shown to increase conjugal recombination frequencies between *E. coli* and *Salmonella* Typhimurium by 1,000-fold (Matic *et al*., 1995), indicating that antisense regulation of *mutL* in *Wolbachia* might explain genome complexity in arthropod strains, which harbour elevated proportions of mobile elements compared to most other obligate intracellular symbionts (Cordaux *et al*., 2008; Kaur *et al*., 2021). The very highly expressed iTSS associated with the *fsa* gene was also unexpected and the lack of enzymatic conserved domain encoded by this transcript is strongly suggestive of an ncRNA-mediated regulatory role. However, a limitation of the current study was that the 3’ end of transcripts are not captured, preventing identification of full-length molecules and restricting structural predictions to the first 100 nt. Nevertheless, detection of TSS associated with ncRNAs reported in previous studies, such as the oTSS linked with *ncrwMel02* (Woolfit *et al*., 2015), provides corroboration that our approach revealed genuine ncRNAs and not biological noise. Furthermore, Woolfit *et al*. (2015) observed upregulation of *ncrwMel02* in testes relative to ovaries in whole flies, suggesting a role in host reproductive manipulation, while we detected downregulation of this transcript under temperature extremes. This may be pertinent to the well-known influence of temperature on the phenotype of CI in *Drosophila* (Clancy and Hoffmann, 1998).

While the genes responsible for CI were identified several years ago (*cifA* and *cifB* in *w*Mel; LePage *et al*., 2017), with homologues referred to as *cidA* and *cidB* in other strains (Beckmann *et al*., 2017), their regulation remains poorly understood. A key question is whether their conserved orientation as a gene pair represents evidence of expression as a bicistronic operon (Beckmann *et al*., 2019; Shropshire *et al*., 2019), which has been challenged by the large disparity in expression levels between the individual genes (*i.e*., an eight-fold higher expression of *cifA* than *cifB* in *Drosophila in vivo*) (Lindsey *et al*., 2018). In the present study, a single pTSS was identified upstream of *cifA* for both *w*MelPop-CLA and *w*AlbB, with no pTSS detected within the intergenic region in *w*MelPop-CLA or upstream of *cifB* in *w*AlbB. This supports a model in which *cifA* and *cifB* can be co-transcribed as a bicistronic transcript as previously determined from RT-PCR evidence, despite the substantial difference in abundance between the individual transcripts. Lindsey *et al*. (2018) identified a Rho-independent transcription terminator within the *w*Mel *cifA-cifB* intergenic region that could provide a mechanism to regulate the two genes if the activity of the terminator is partial. Other potential mechanisms that could reconcile the data supporting co-transcription and differential regulation between *cifA* and *cifB* are the presence of a Puf family-like RNA-binding domain within the *cifA* gene (Lindsey *et al*., 2018) and/or the use of alternative TSS. Although expression of the *cifB* asTSS in *w*AlbB was weak under the conditions tested here and not conserved in *w*MelPop-CLA, it highlights the complexity of regulation of operons and the possibility of “fine tuning” of *cifA* and *cifB* expression, which in whole organisms may be temporal, tissue-specific, or calibrated by host sex.

Since the late 1990s, extensive efforts have been made to develop antibiotic regimens that can safely eliminate adult filarial worms in onchocerciasis and lymphatic filariasis by targeting their *Wolbachia* symbionts, which in contrast with most arthropod *Wolbachia* strains, are essential for the viability and reproductive physiology of their hosts (Johnston *et al*., 2021). Initially, doxycycline (a tetracycline derivative) garnered the most interest but regimens of several weeks are required for killing of adult worms, which led to evaluation of antibiotics from a variety of other classes (*e.g*., rifamycins, fluoroquinolones and macrolides), some of which clear *Wolbachia* more rapidly than doxycycline in preclinical assays (Johnston *et al*., 2014). As no stable culture system for filarial endosymbionts has been established, arthropod *Wolbachia* strains (especially *w*AlbB) in mosquito cell lines have been utilized as a substitute (Hermans *et al*., 2001; Johnston *et al*., 2014).

Very few studies have attempted to characterise the global response of *Wolbachia* to antibiotic stress, especially in arthropod strains. However, we previously performed RNA-Seq in a TEX experiment (combined with proteomics) on *w*MelPop-CLA following exposure to doxycycline *in vitro* for three days (Darby *et al*., 2014). Many similarities were apparent between wMelPop-CLA in the TEX study and the current Cappable-Seq study of *w*AlbB after 24 hr exposure to doxycycline. These included downregulation of HSP genes; and upregulation of ribosomal protein genes, enzymes involved in nucleotide or lipid metabolism, the twin-arginine translocation (Tat) system, and transposase genes. There were also a few differences, such as upregulation of some outer membrane protein (OMP) genes and an ankyrin (both downregulated in *w*MelPop-CLA), whereas genes of the T4SS were upregulated in *w*AlbB but unaffected by doxycycline in *w*MelPop-CLA. Importantly, this does not necessarily indicate dissimilar responses between strains, as both the primary transcript enrichment method (TEX *v*. Cappable-Seq) and the antibiotic exposure period (one or three days) are likely to have had a considerable impact on the comparability of the datasets.

While there were distinctive TSS expression profiles and DE associated with antibiotic stress compared with temperature stress, there were also areas of overlap, such as downregulation of HSP genes following cold stress or exposure to doxycycline. Indeed, responses to antibiotics are known to stem from conserved bacterial stress response networks, with tetracyclines inducing a cold-shock response in *E. coli* that includes downregulation of *rpoH* (though some HSP genes are nevertheless upregulated), as well as synergistic effects on *E. coli* viability when combined with cold stress (Cruz-Loya *et al*., 2019). Doxycycline-induced ribosomal stalling produces incompletely synthesized polypeptides that are then recycled via protease activity (Keiler, 2015). Doxycycline exposure induced upregulation of a gene encoding an inner membrane-bound protease, *ftsH*, in *w*AlbB, which contributes to the degradation of proteins for amino acid (AA) recycling (Langklotz *et al*., 2012). FtsH-mediated proteolysis is also involved in the post-translational regulation of the heat-shock regulator RpoH (Tomoyasu *et al*., 1995). Thus, proteolytic degradation of RpoH by FtsH could contribute to downregulation of heat-shock genes whilst providing an AA pool for nutritional uptake.

Changes to the structure of the Gram-negative outer membrane and activity of transporters spanning the periplasmic space are common during exposure to antibiotics, which can be accompanied by alterations to phospholipid metabolism. A pTSS associated with the *plsC* gene (DEJ70_RS02625), which encodes 1-acyl-sn-glycerol-3-phosphate acyltransferase of the glycophospholipid metabolism pathway, was the most upregulated pTSS in response to doxycycline exposure. PlsC is an integral membrane protein responsible for the catalytic conversion of 1-acyl-sn-glycerol-3-phosphate into 1,2-diacyl-sn-glycerol 3-phosphate (phosphatidic acid, PA) (Coleman, 1990), which in turn is the major precursor for membrane phospholipids such as phosphatidylethanolamine, phosphatidylglycerol, and cardiolipin. In parallel, upregulation of genes encoding a major facilitator superfamily (MFS) transporter, a resistance-nodulation-cell division (RND) permease subunit, and the outer membrane efflux protein TolC were observed. Both MFS transporters and the efflux pumps of the RND superfamily are associated with multidrug resistance in *E. coli* and other Gram-negative bacteria (Fernando and Kumar, 2013; Greene *et al*., 2018; Kumar *et al*., 2020). TolC is a component of tripartite efflux pumps, including the RND superfamily, MFS transporters, and the type I secretion system. In the latter, HlyD (also upregulated in response to doxycycline) is an adaptor that facilitates transport of large proteins across the periplasmic space (Greene *et al*., 2018). In *E. coli*, mutations in *tolC* led to a five to six-fold decrease in tetracycline resistance conferred by Tet(A) pumps, which this was demonstrated to result from disruption to an RND efflux system (de Cristóbal *et al*., 2006). Only a single gene associated with secretion or transport, *yidC*, was downregulated after doxycycline transport, which encodes a membrane protein insertase. Notably, in *Aeromonas hydrophila,* knockout of *yidC* substantially lowered susceptibility to several antibiotics of the tetracycline class (Yao *et al*., 2018).

Further evidence for membrane remodelling following doxycycline exposure was apparent from upregulation of pTSS associated with OMPs such as peptidoglycan-associated lipoprotein (PAL), a member of the OmpA-like family and the primary pathogen-associated molecular pattern in *Wolbachia* cells (Turner *et al*., 2009). This OMP has been partially characterized in *E. coli* and other Gram-negative pathogens such as *Pseudomonas aeruginosa* (where the PAL homologue is OprL) and *Haemophilus influenzae* (homologue is P6). Close associations between PAL and OmpA itself (or its homologues) confer integrity to the outer membrane (Navare *et al*., 2015), and mutants lacking P6 in *H. influenzae* have greatly increased susceptibility to antibiotics (Murphy *et al*., 2006). Few RNA-Seq studies have been performed on Gram-negative bacteria exposed to doxycycline, but in *Yersinia pseudotuberculosis*, two OMP genes (including a lipoprotein gene, although not a PAL homologue) were the only genes found to be upregulated following treatment and were associated with doxycycline tolerance (Alvarez-Manzo *et al*., 2022).

Rifampicin (a rifamycin targeting RNAP) or moxifloxacin (a fluoroquinolone targeting DNA gyrase and topoisomerase IV) had a much more limited impact on DE of TSS in the 24-hr timescale of our experiments compared with doxycycline. Over the past 25 years, the susceptibility of *Wolbachia* to various antibiotics (including rifampicin and moxifloxacin) has been established *in vitro* by adding compounds to mosquito cell lines infected with insect-derived *Wolbachia* strains, or by incubation of adult filarial worms in culture media containing drugs (Johnston *et al*., 2014). Cell line-based studies have differed between laboratories in whether *Wolbachia* are inoculated into host cells simultaneously with, or prior to, drug exposure, which leads to different growth kinetics (Johnston *et al*., 2014). In the Aa23 cell line, the doubling time of *w*AlbB has been estimated as 14 hr (Fenollar *et* al., 2003), while in adult filarial worms, turnover of symbiont populations is very slow except in the female reproductive system (McGarry *et al*., 2004). Antibiotic susceptibility testing is performed over a period of at least several days, sometimes with replenishment of drugs midway through the assay (Johnston *et al*., 2014). In our study, we harvested cells 24 hr after antibiotic exposure, since a longer treatment period would have increased the proportion of dead bacteria and reduced *Wolbachia* RNA yields. Similarly, host cells were sub-cultured several days prior to treatment to ensure that *Wolbachia* density had plateaued at a level permitting recovery of sufficient symbiont RNA for high transcriptome sequence coverage. In contrast, RNA-Seq experiments on free-living bacteria exposed to fluoroquinolones added the antibiotic to liquid medium during mid-late exponential phase (Ulrich *et al*., 2013; Sinel *et al*., 2017). This may explain why by 24 hr post-treatment, we did not observe any signs of the global SOS response in *Wolbachia* exposed to moxifloxacin, whereas SOS induction is typical in free-living bacteria following short-term incubation with fluoroquinolones.

Although the number of TSS exhibiting DE after rifampicin treatment was small, it was clear that this drug had an opposite effect to doxycycline on genes involved in secretion and transport. Rifampicin triggered downregulation of T4SS components and an RND transporter permease (both upregulated by doxycycline), whereas *yidC* was upregulated by rifampicin but downregulated by doxycycline. These results underscore specific aspects of regulation in *Wolbachia* following exposure to stress within a larger framework of stress-responsive genes. Studies in *E. coli* have shown that while rifampicin blocks the activity of free RNAP molecules, those already bound to DNA and participating in transcription are not inhibited, permitting residual RNA synthesis (Chen *et al*., 2015). Thus, the lack of broadscale downregulation of *Wolbachia* transcription by 24 hr observed here is compatible with persistent RNAP activity at TSS across the genome. Experiments designed to characterise the non-culturable state in *Mycobacterium tuberculosis* use potassium ion depletion to induce a dormant state followed by rifampicin treatment, which kills dividing bacilli but not dormant persisters. However, ongoing transcriptional activity in dormant *M. tuberculosis* is thought to result from greatly extended transcript stability rather than ongoing transcription *de novo* under drug pressure (Ignatov *et al*., 2015). Notably, in long-term experiments in animal models of filariasis, rifampicin failed to completely clear *Wolbachia* symbionts (Bah *et al*., 2014), leading to recrudescence from persisting populations in the reproductive tract of female worms in the absence of mutations in the *Wolbachia* genome (Gunderson *et al*., 2020). Hence, these filarial *Wolbachia* displayed rifampicin tolerance rather than resistance. One candidate molecule that might contribute to persistence is RNase P, a ribozyme that is essential for the processing of precursor tRNA transcripts, which was upregulated after rifampicin exposure in *w*AlbB. Accordingly, an increase in RNase P levels after rifampicin treatment has also been observed in *Bacillus subtilis* (Dittmar *et al*., 2004).

### Conclusions

This study detected statistically significant DE amongst all known TSS types for both *w*MelPop-CLA and *w*AlbB. This highlights the severe limitations of conventional RNA-Seq, in which distinct transcriptional units are merged and quantified in aggregate. Moreover, it suggests that the characterisation of *Wolbachia* (and other obligate intracellular bacteria) as organisms with limited capacity for transcriptional regulation is premature (Chung *et al*., 2020), reflecting technology and not biology. At a minimum, the sophisticated reproductive manipulations induced by arthropod *Wolbachia* strains are likely to be stringently controlled between host tissues and sensitive to host sex (Gutzwiller *et al*., 2015; Harumoto and Fukatsu, 2022). However, the transcriptional landscape should also be revisited in the filarial symbionts with more reduced genomes to determine their TSS repertoire and potential for regulation across the lifecycle, and/or between the somatic and germline cells of the host (Darby *et al*., 2012; Chevignon *et al*., 2021). This is particularly important for understanding transcriptional plasticity under drug pressure (Darby *et al*., 2014) and its relationship to the challenges of eliminating *Wolbachia* with short antibiotic regimens for the control of filarial diseases.

Our data show that the differences in thermal susceptibility observed between *w*AlbB and *w*MelPop-CLA (Ross *et al*., 2017) are not due to a failure by the latter to induce a heat-shock response; indeed, only wMelPop-CLA displayed regulation of *rpoH* following temperature stress. Instead, differences in host background, specifically hosts that share a longer phylogenetic history with their symbiont, may impact host-symbiont homeostasis during stress conditions. While the *wspB* polymorphisms previously identified (Gu *et al*., 2022; Hague *et al*., 2022; Martinez *et al*., 2022) play a key role in thermal sensitivity, future studies should focus on how the differential regulation of ncRNAs and transposase activation impact the resilience of *Wolbachia* strains exposed to heat and other stressors. Applying Cappable-Seq to *Wolbachia* in whole invertebrate host systems will provide further insights into this remarkably adaptable and successful symbiont.

## MATERIALS AND METHODS

### Cell Culture

To investigate the transcriptional response of *w*MelPop-CLA and *w*AlbB exposed to stress, mosquito cell lines were used as a model to study *Wolbachia* gene expression within the background of a host relevant to current vector control efforts. The RML-12 and Aa23 *Ae. albopictus* cell lines harbouring *w*MelPop-CLA (McMeniman *et al*., 2008) and *w*AlbB (O’Neill *et al*., 1997) respectively were maintained at 28°C in T75 flasks. Cell lines were grown in 15 ml insect media composed of 45% Mitsuhashi & Maramorosch medium (HiMedia), 45% Schneider’s *Drosophila* medium (Sigma), 10% foetal calf serum, and 0.18% L-glutamine. Cell lines were passaged every 4-6 days until confluency in a 1:5 ratio (3 ml suspended cells plus 12 ml fresh media).

### Temperature stress induction

For temperature stress, mosquito cells were maintained in either T25 (RML-12: 5 ml cultures) or T225 (Aa23: 45 ml cultures) flasks at 28°C for 5 - 7 days and then transferred to incubators at each temperature (21°C, 28°C, 34°C) for either 4, 24, or 48 hr before harvest, with three biological replicates per condition. Temperatures were selected to reflect values experienced in tropical climates such as the wet season in Cairns, Australia (Richardson *et al*., 2013) whilst avoiding temperature limits that may damage the host cell line during continuous exposure (Dobson and Rattanadechakul, 2001). Once the incubation periods were complete, cells were prepared for RNA extraction.

Due to the presence of low-density *w*MelPop-CLA, RML-12 sample preparation differed to Aa23 as follows. The RML-12 cells were processed by resuspension and transfer into 50 ml Falcon tubes placed directly onto ice to stall metabolic and transcriptional activity. Five ml of sterile 3 mm glass beads were added to the cell suspension and vortexed for 5 minutes to lyse mosquito cells and liberate intracellular *Wolbachia*. The vortexed cell suspension was then centrifuged at 2,500 × g at 4°C for 15 min to pellet host cell debris. The *Wolbachia*-containing supernatant was subsequently filtered through a 5 μm syringe filter then further centrifuged at 18,000 × g at 4°C for 15 minutes to pellet *Wolbachia* cells. The supernatant was removed and the *Wolbachia* pellet was resuspended in 1 ml media and further filtered through a 2.7 μm syringe filter into a 1.5 ml Eppendorf tube. After a final centrifugation step at 18,000 × g at 4°C for 15 min, the *Wolbachia* pellets were suspended in 1 ml QIAzol and stored at -80°C. For Aa23, cells were resuspended and transferred into 50 ml Falcon tubes and centrifuged at 300 *g* at 4°C for 10 minutes. Aa23 cells were also resuspended in and transferred to 50 ml Falcon tubes but did not undergo cell bead lysis; instead the cell suspension was centrifuged at 2,500 × g at 4°C for 15 min followed by removal of the supernatant. The remaining cell pellet was then suspended in 1 ml QIAzol and stored at -80°C.

### Antibiotic stress induction

The *w*AlbB antibiotic stress induction was performed by growing Aa23 cells in T25 flasks (5 ml culture, triplicate) for six days. Antibiotics were then added at 0.25 μg/ml (doxycycline), 0.25 μg/ml (rifampicin), and 1 μg/ml (moxifloxacin) for 24 hr. Cells were then harvested by resuspending cells and transferring them into 15 ml Falcon tubes. Samples were centrifuged at 300 x g for 5 minutes, the supernatant removed, and 2 ml of QIAzol added to each cell pellet prior to storage at -80°C. Antibiotic dosages were based on physiologically relevant concentrations and those chosen by previous studies (Fenollar, Maurin and Raoult, 2003; Darby *et al*., 2014; Johnston *et al*., 2014).

### RNA extraction

RNA was extracted via QIAzol-chloroform extraction. Briefly, 200 μl of 100% chloroform was added per millilitre of QIAzol-suspended sample and shaken by hand for 15 seconds. Samples were incubated at room temperature for three minutes and then centrifuged at 12,000 *g* at 4°C for 15 minutes. After incubation, 500 μl of the aqueous phase was transferred to DNA LoBind tubes (Eppendorf) to which 750 μl of absolute ethanol was added. Total RNA was extracted from each samples using the Monarch total RNA miniprep kit (NEB product number T2010S) TRIzol extraction protocol with on-column DNase treatment, as specified by the manufacturer. The RNA concentrations were assessed using a NanoDrop spectrophotometer (ThermoFisher Scientific).

### Cappable-Seq enrichment

For the capping reaction, 30 μl of 2 - 10 μg of DNase treated RNA was added to 5μl of 10X vaccinia capping enzyme buffer (NEB product number M2080) with 5 μl of 5 mM 3’ -desthiobiotin-guanosine triphosphate (DTB-GTP) (NEB product number N0761), 5 μl vaccinia capping enzyme (NEB product number M2080), 5 μl inorganic pyrophosphate (New England Biolabs product number M2403), and ultrapure water (as needed) to a total volume of 50 μl and incubated at 37°C for 1 hr. Samples then underwent RNA purification using the Monarch RNA clean-up kit (NEB product number T2030S), modified to increase the washing step to a total of five times to ensure removal of free DTB-GTP; RNA was then transferred to a PCR tube. The DTB-GTP capped RNA then underwent fragmentation by adding 1.25 μl of 10X T4 polynucleotide kinase buffer and was incubated at 94°C for 5 minutes, then placed directly on ice. The removal of the 3’ phosphates from fragmented RNA was accomplished by adding 7 μl of 10X T4 polynucleotide buffer and 3.2 μl of ATP-free T4 polynucleotide kinase, followed by incubation at 37°C for 15 minutes. Samples then underwent RNA purification using the Monarch RNA clean-up kit (NEB T2030S) and were eluted with 40 μl ultrapure water.

Samples were subjected to two rounds of enrichment with streptavidin beads by adding 30 μl of DTB-GTP capped RNA to 30 μl of preprepared streptavidin beads and placed onto a rotation mixer at room temperature for 30 minutes. Samples were then placed onto a magnetic stand for 5 minutes or until the solution was clear. Whilst on the magnetic stand, samples were washed four times with 200 μl washing buffer (10 mM Tris-HCL pH 7.5, 120 mM NaCl, 1 mM EDTA). Beads were resuspended in 11 μl low TE elution buffer and incubated in a thermomixer at 80°C for 10 minutes before a magnetic stand was used to facilitate recovery of RNA. Bead resuspension was performed twice before a second round of streptavidin enrichment was performed with 20 μl of DTB-GTP capped RNA and 20 μl of streptavidin beads. Decapping of the 5’ DTB-GTP cap was performed to leave the 5’ monophosphate for ligation by adding 20 μl of capped RNA to 2.2 μl of 10X ThermoPol buffer (NEB product number B9004) and 2 μl RppH (NEB product number M0356S), followed by incubation at 37°C for 1 hr. Next, 1 μl of 1:10 diluted proteinase K was added to the solution and incubated at 37°C for 10 minutes. The RNA solution then underwent a final round of RNA purification using the Monarch RNA clean-up kit (NEB T2030S) before library preparation.

### Library preparation and sequencing

Enriched RNA underwent library preparation using the NEBNext multiplex small RNA library prep kit (NEB product number E7300S) following the manufacturer’s instructions. After library completion, samples underwent PCR clean up by adding 1 volume of Ampure beads followed by incubation at room temperature for 10 minutes. Samples were placed onto a magnetic stand until the solution became clear. The supernatant was removed and beads were washed with 80% ethanol four times, air-dried, resuspended in 40 μl low TE buffer, and incubated at room temperature for 10 minutes. The supernatant was collected, then the resulting library was diluted 1:10 and analysed using the high sensitivity DNA Agilent Bioanalyzer protocol using a DNA 1000 chip according to the manufacturer’s instructions. Samples were sequenced on an Illumina NEXTSeq using high output single-end 150 bp sequencing.

### Mapping

Adapter trimming and contaminant filtering was performed using BBduk from the BBtools software package (version 38.86). Unpaired single-end reads were mapped using Bowtie2 (version 2.4.5) with the local alignment option. Reads were mapped simultaneously onto a concatenated fasta file containing the genomes of *Ae. albopictus* mitochondria (accession NC_006817), *Ae. albopictus* (assembly GCA_001876365.2), and *w*Mel (assembly GCA_000008025.1) or *w*AlbB (assembly GCA_004171285.1). A threshold for a minimal sequencing depth (supplemental FS4) was chosen based on the recommended minimal sequencing depth of two million for differential gene expression in *E. coli* (Haas *et al*., 2012). The two million *E. coli* minimal read depth equates to 0.43 million reads per Mb of genome length, or a minimal desired library depth of 0.54 M and 0.63 M reads for *w*MelPop-CLA and *w*AlbB, respectively. The *w*MelPop-CLA genome is highly similar to *w*Mel, besides defined minor differences (Woolfit *et al*., 2013), allowing *w*Mel to be a suitable proxy for gene expression in wMelPop-CLA. Aligned reads that mapped to *Wolbachia* were further filtered for those with a unique mapping score of mapq20 using samtools (version 1.11) to ensure reads associated with repetitive regions were confidently assigned. Removing non-uniquely mapped reads may limit additional information from repeat elements, such as transposons; however, their removal provided greater confidence in expression of TSS that remained associated with repeat elements. Reads were then converted into count values using the software HTSeq-count (version 0.11.1).

### TSS Calling

TSS calling of mapped reads were conducted using the TSS calling software pipeline (accessible from https://github.com/Ettwiller/TSS) using a clustering filter of a 10-nucleotide window. Designating a TSS type was performed using a custom python script implementing pandas and Bedtools closest (v2.29.2). The TSS types were assigned based on location of the TSS relative to closest neighbouring genes using the following criteria: pTSS had the highest read count within 300 nucleotides upstream of the start codon; gTSS were within 300 nucleotides upstream to the start codon, but with read counts less than the pTSS; iTSS were internal to the CDS; asTSS were antisense to coding sequence and/or 100 nucleotides downstream to the antisense stop codon; and oTSS comprised those TSS not fulfilling any of the previous definitions. High confidence TSS were determined by only selecting those TSS which were present in all replicates within a condition. Any TSS that were absent in a condition but present in others were given a prior count of 10. To apply the same minimal expression value to all samples, the threshold for a TSS to be called was 10 CPM, based on the smallest sample library for temperature and antibiotic experiments separately. The TSS statistics and graphs were generated via the python library packages matplotlib and seaborn. Heatmaps were generated using heatmap.2 in the R package ggplot2 (version 3.3.2).

### Orthologous gene comparisons

To define orthologous genes between *w*Mel and *w*AlbB, the amino acid sequences of all protein-coding genes in each genome were compared via the OrthoFinder package (Version 2.5.4). As a conservative approach to increase the likelihood of functional similarity between protein-coding genes, only those genes with protein products that shared one-to-one homology between *w*Mel and *w*AlbB were considered for further analysis.

### Motif discovery

To find sequence motifs indicative of promoter regions, the upstream 50-nucleotide regions of all TSS were used as input into the web browser-based MEME suite (Version 5.4.1) with up to 10 motifs patterns selected in the settings. Motifs were visualised using the software package WebLogo (Version 2.8.2).

### Differential expression analysis

Differential expression (DE) analysis was conducted using the empirical analysis of digital gene expression package edgeR (version 3.30.3) with the following settings: prior count of two, minimal fold change threshold of two, and an adjusted p-value false discovery rate (Benjamini-Hochberg procedure) of 5%. Multidimensional scaling plots were analysed and produced via the plotMDS function within edgeR.

### Functional annotation

To aid in the biological interpretation of DE TSS under stress, genes were assigned Kegg Ortholog (KO) annotations so they could be included in the Kyoto Encyclopedia of Genes and Genomes (KEGG) pathway analysis. Functional gene annotations were retrieved with the use of EggNOG (version 5.0.0); KO annotations of DE genes were then used as input into the KEGG pathway analysis under organism prefixes of ‘wol’ and ‘wpp’ for *w*Mel and *w*AlbB, respectively. Pathways which resulted in the greatest number hits or were associated with highly DE TSS were the focus of biological interpretation.

### RNA secondary structure analysis

To evaluate the likelihood of whether a TSS could be associated with an ncRNA instead of, or associated with, a protein-coding gene annotation, the 100 nucleotide downstream sequences of all TSS types were analysed for predicted RNA secondary structures with MFE values in RNAfold (version 2.4.13) of the ViennaRNA package (version 2.0) using default settings. A total of 2,000 (1,000 per strand) randomly generated TSS locations were created with a custom python3 script as a comparative control.

### Operon structure analysis

To assess the structure of predicted operons within *w*MelPop-CLA and *w*AlbB, gene annotations for both strains were used as input for the Rockhopper tool (version 1.7.0.11). The BAM alignment files from 28°C at 48 hr conditions were used as input into Rockhopper to decipher whether the intergenic gaps between genes displayed evidence of co-expression. Operons and associated TSS were visualised using the Artemis genome browser and annotation tool (version 18.1.0). Expression of an operon was positively defined if the initiating gene of a predicted operon harboured a pTSS. Operons were considered to contain an ATU if they displayed a moderately expressed intergenic TSS upstream to a gene within the associated operon.

## Supporting information

Combined supplement

## Acknowledgements

We gratefully acknowledge the World Mosquito Program for supply of the RML-12 cell line infected with *w*MelPop-CLA. We thank Laurence Ettwiller, Ira Schildkraut and George Tzertzinis for helpful discussions and expertise on the Cappable-Seq protocol.

## Data Availability

Raw data are available at the SRA database under BioProject accession no. PRJNA1145481.

## Notes

### Competing Interest Statement

The authors have declared no competing interest.

