## Supplementary material for "Cappable-Seq reveals the transcriptional landscape of stress responses in the bacterial endosymbiont *Wolbachia*": Combined supplement

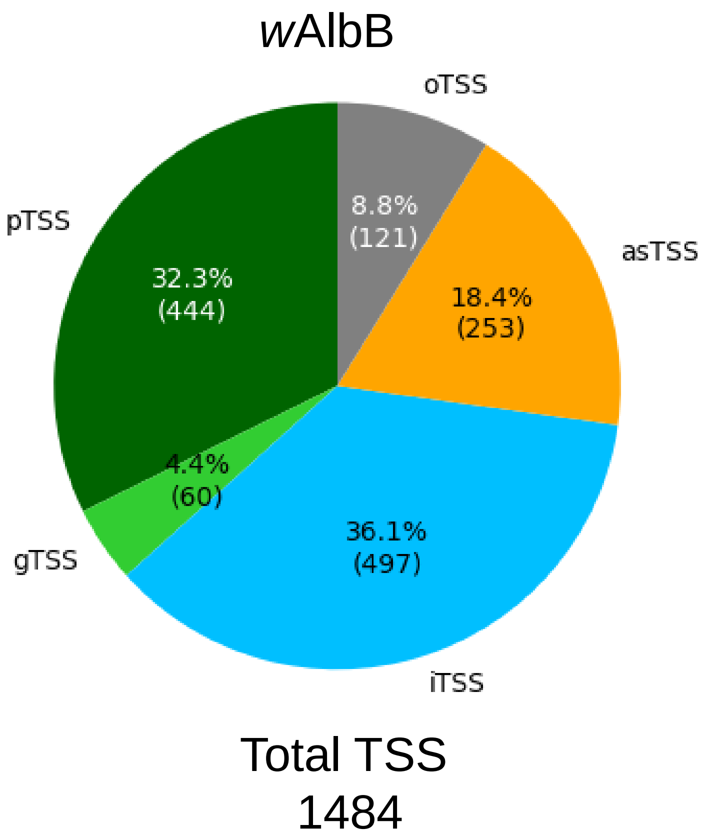


**Figure S1.** Summary of wAlbB TSS types across all conditions tested using the CPM threshold identical to *w*MelPop-CLA (25.48 CPM), representing the CPM of 10 reads in the smallest library that passed the recommended depth (0.54 M and 0.63 M reads for *w*MelPop-CLA and *w*AlbB respectively).

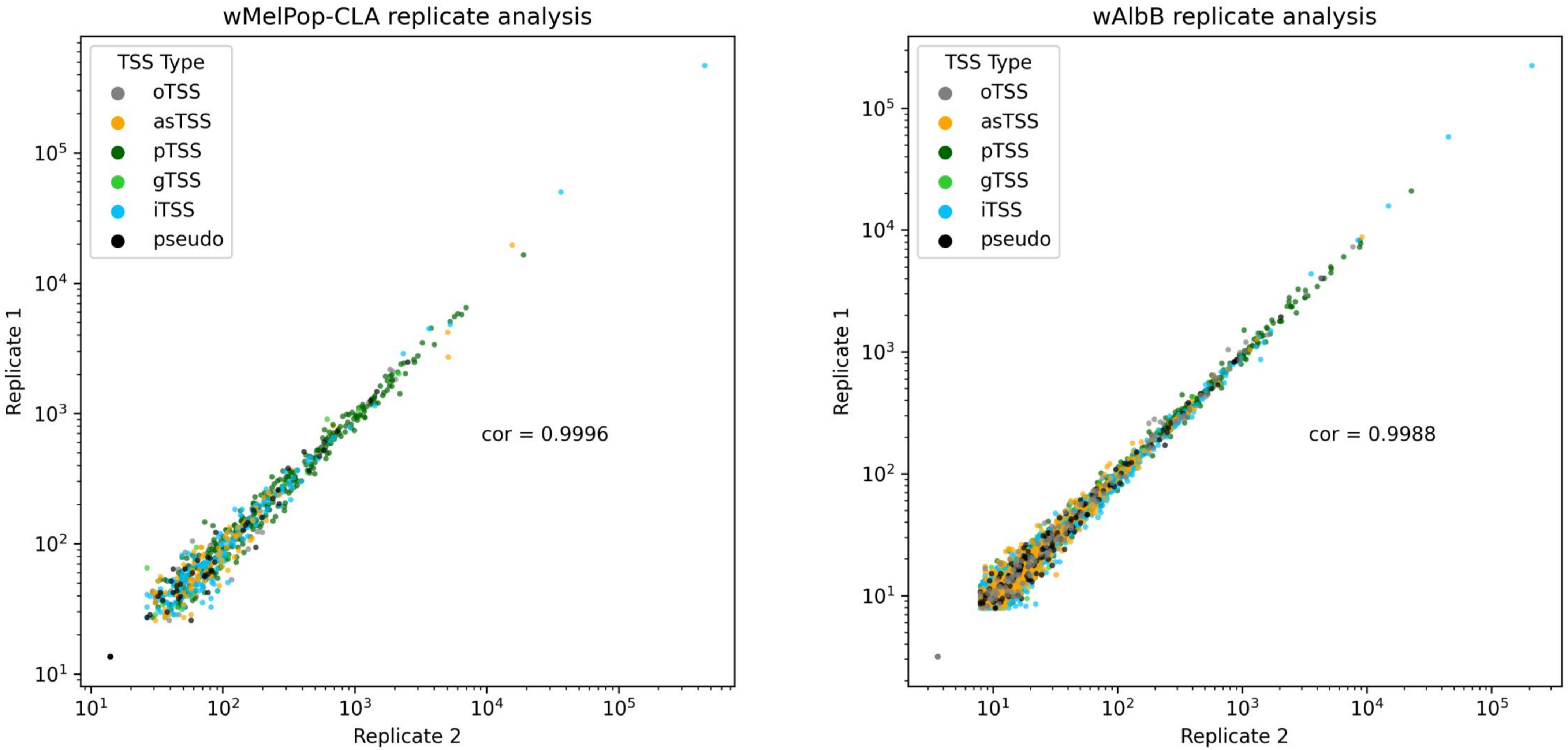


**Figure S2**. Cappable-Seq expression between replicates. Scatter graphs of TSS expression between 28℃ replicates of A) *w*MelPop-CLA and B) *w*AlbB. Each dot represents the CPM of each TSS coloured by its designated TSS type. Cor = correlation coefficient.

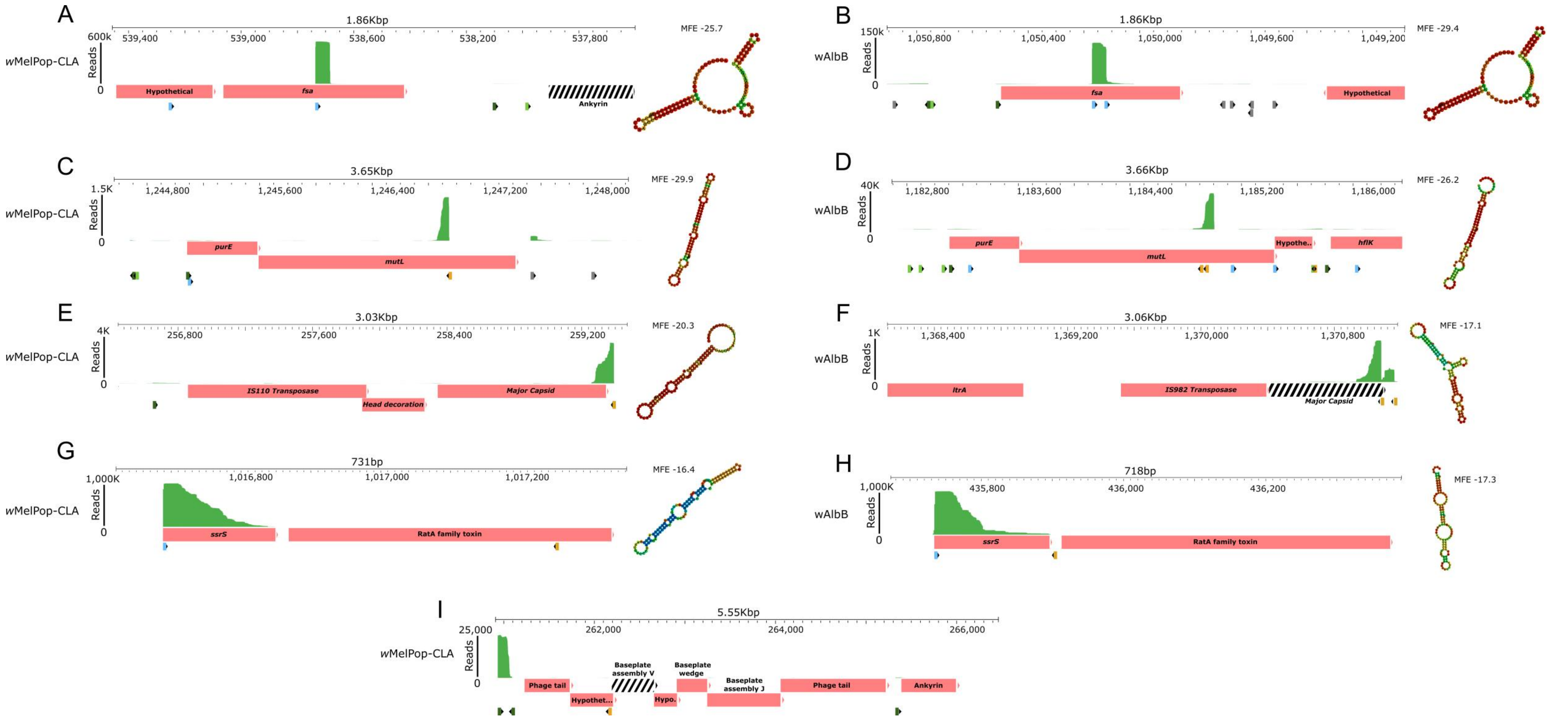


**Figure S3.** Overview of TSS in select regions of interest. The iTSS of *fsa* for **A** *w*MelPop-CLA and **B** *w*AlbB. The asTSS of *mutL* for **C** *w*MelPop-CLA and **D** *w*AlbB. The asTSS of a major capsid gene for **E** wMelPop-CLA and **F** *w*AlbB. The iTSS of *ssrS* (6S) for **G** *w*MelPop-CLA and **H** *w*AlbB. The pTSS of a phage tail gene in **I** wMelPop-CLA. Genes are coloured red boxes, pseudogenes are black and white striped. Green peaks represent mapped reads under 28°C. The TSS are placed below associated genes and coloured per TSS type as dark green, pTSS; light green, gTSS; light blue, iTSS; orange, asTSS; grey, oTSS. Predicted RNA secondary structures for the 100 nt downstream sequence are placed to the right of each displayed region along with the associated MFE value.

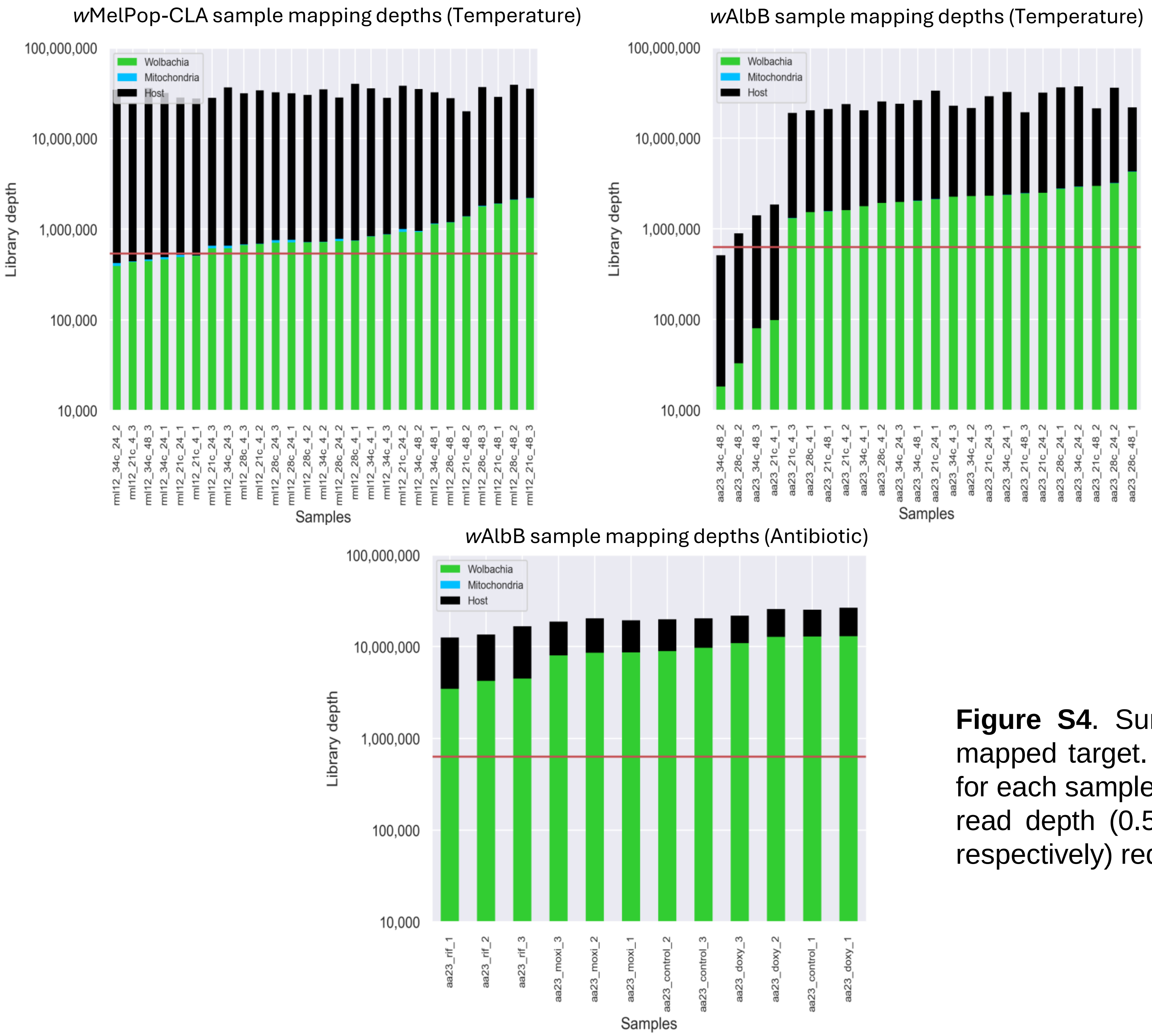


**Figure S4**. Summary of mapped reads across conditions and their mapped target. Bar chart representing reads depths of mapped reads for each sample. Red horizontal line represents the suggested minimum read depth (0.54 M and 0.63 M reads for *w*MelPop-CLA and *w*AlbB respectively) required for robust DE analysis.

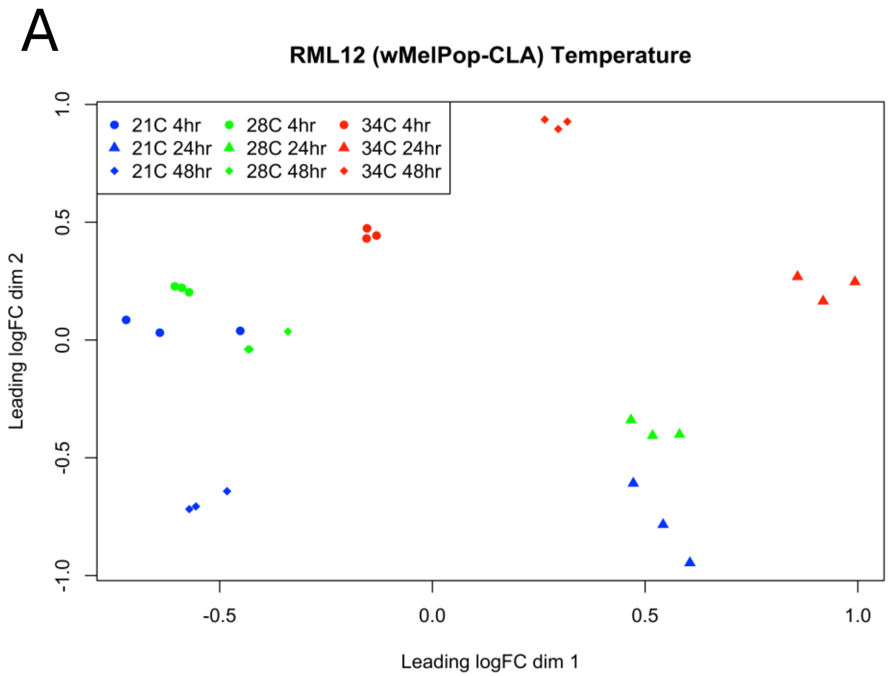


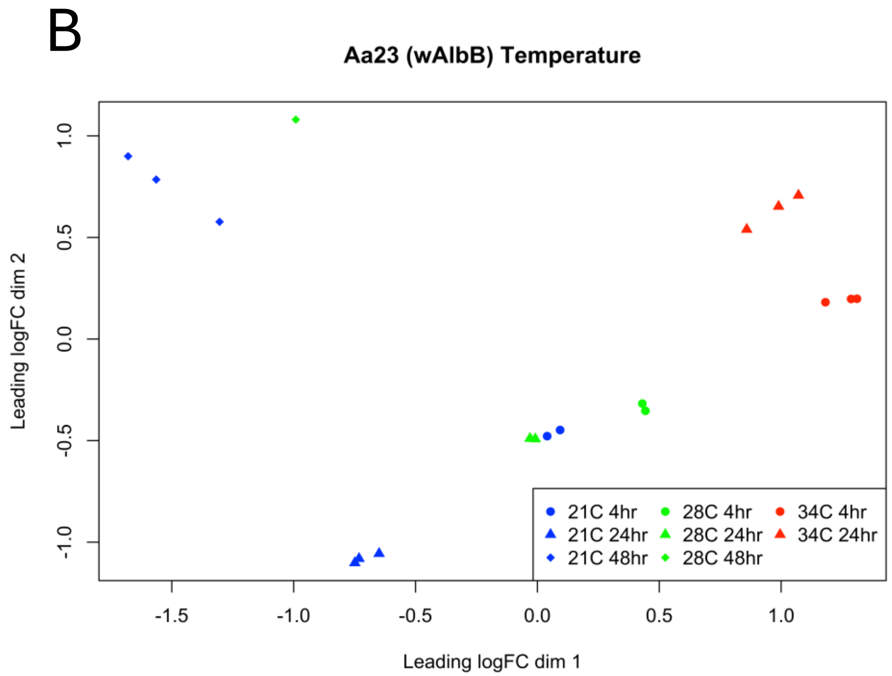


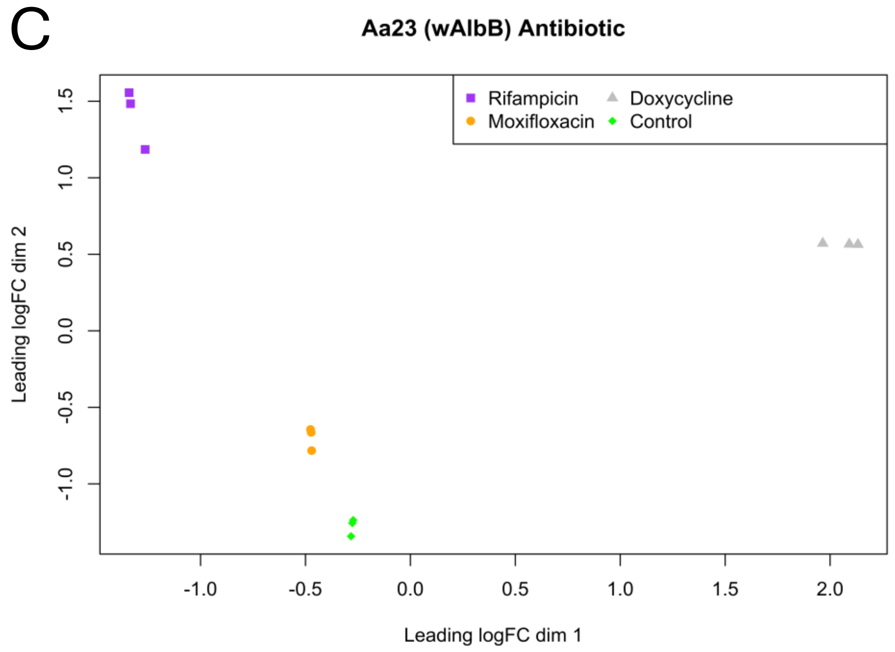


**Figure S5**. MDS plots of stress-induced *w*MelPop-CLA and *w*AlbB samples. A) *w*Mel-PopCLA exposed to temperature stress, B) *w*AlbB exposed to temperature stress, C) *w*AlbB 24 hr exposure to antibiotic stress.

**Table S1.** Summary of TSS-assisted operon prediction. Occurrence of predicted operons with number of associated genes for both *w*Mel and *w*AlbB genomes.

| Genes in operon | wMel | wAlbB |
|---|---|---|
| 28 | 1 | |
| 23 | | 1 |
| 10 | 1 | |
| 9 | | 1 |
| 8 | 1 | |
| 7 | 4 | 2 |
| 6 | 3 | 4 |
| 5 | 8 | 2 |
| 4 | 20 | 21 |
| 3 | 32 | 32 |
| 2 | 163 | 149 |
| Total | 233 | 212 |

**Table S2.** TSS types associated with TSS-assisted operon prediction. Summary of operons with associated TSS types for both *w*Mel and *w*AlbB genomes.

| | Operons | pTSS | gTSS | iTSS | asTSS |
|---|---|---|---|---|---|
| wMel | 233 | 40 | 2 | 24 | 13 |
| wAlbB | 212 | 58 | 10 | 84 | 56 |
